# Internalized Components of Membrane Attack Complexes Disrupt Proteostasis and Acquire Alarmin-Like Properties

**DOI:** 10.64898/2026.07.03.736247

**Authors:** Guiyu Song, Zihan Ma, Matthew Fan, Liying He, Yulong Lan, Weihao Li, Zhengtao Jiang, Quan Jiang, Dylan P. Noone, Andrea Nans, Kamal L. Nahas, Mahsa Nouri Barkestani, Shaoxun Wang, Qianxun Wang, Pengwei Ren, Jolin Cheng, Yinuo Zang, Haitian Zhou, Justin Johnson, Clancy Mullan, Xiangyu Gong, Doryen Bubeck, Gilbert Moeckel, Michael Mak, George Tellides, Dan Jane-wit

## Abstract

Immune effects of membrane attack complexes (MAC) have been widely attributed to their abilities to cause cell death. Here, we show that the MAC component, C9, forms non-cytolytic aggregates with pro-inflammatory effects. Intracellular aggregates of C9 are detected within inflamed tissues of patients in association with endothelial cell (EC) activation but not increased cell death. We identify NUMBL as a Rab35 effector that directly binds surface-bound C9 to promote C9 internalization and entry into the endolysosomal pathway. Within acidified endolysosomes, C9 forms insoluble aggregates that are targeted for degradative aggrephagy in a process that activates NF-κB. For C9 aggrephagy to occur, ZFYVE21, a Rab5 effector, complexes with RNF34 to bridge C9 aggregates to LC3B+ aggresome membranes. We detect C9 aggregates *in vivo*, and we show that a ZFYVE21-RNF34 signaling axis is required for C9 aggrephagy and NF-κB -dependent EC activation in three separate mouse models. Mice with conditional loss of ZFYVE21 in ECs show reduced aggregraphy, resulting in attenuated systemic inflammation and reduced tissue injury following skin transplantation. Our data show that the C9 component of MACs forms intracellular aggregates with alarmin-like properties.

## INTRODUCTION

Membrane attack complexes (MAC) are immune mediators promoting tissue injury^1^ that have been therapeutically targeted in ∼40 inflammatory conditions.^2^ Among these, in antibody-mediated rejection (ABMR), complement proteins inclusive of MACs show diagnostic,^3^ prognostic,^4^ and therapeutic value.^2, 5, 6^ In C4d^+^ ABMR, MACs principally assemble on endothelial cells (EC) in association with dysregulated inflammation but not diffuse EC death.^7, 8, 9^A similar process is observed in connective tissue disorders,^10^ autoimmune vasculitis,^11^ and viral infections including SARS-CoV-2,^12, 13^ suggesting that MACs show a generalizable form of non-cytolytic immunogenicity.

Whilst promoting osmolysis of bacterial xenogens and anucleated red blood cells, MACs show complex mechanism(s) when deposited on autologous, nucleated cells like ECs.^1, 14^ MAC deposition on ECs triggers membrane remodeling events where MAC-related components can either be shed by plasma membrane blebbing or internalized via clathrin-dependent^15^ and clathrin-independent endocytosis.^16^ MACs may further exert non-cytolytic or sublytic effects, activating signaling related to cell stress, proliferation, and survival.^17, 18, 19^

In the setting of ABMR, MACs generate inflammatory signals that promote EC activation *in vitro* and *in vivo* without increased cell death.^20^ Here, MAC components are internalized and trafficked to Rab5^+^ endosomes^15, 21^ that subsequently employ effector molecules^22, 23^ to activate NF-κB.^15, 21, 22, 23, 24, 25^ Attendant to this process is the fact that MAC pores are large, consisting of ∼ 20–25 complement proteins and with molecular weights of ∼ 1,600-2,000 kD.^1, 5^ Upon their internalization, the large, *de novo* protein burden of MAC-associated components is expected to impose significant challenges to proteostasis, particularly during chronic ABMR (CABMR) where MACs may become persistently assembled and internalized on a given EC over time. How ECs cope with the large burden of internalized MAC proteins is unclear, and molecular coping mechanisms are undefined.

The intracellular role of complement is an emerging area in which immune activation mechanisms are interlinked with trafficking processes regulating homeostasis. Intracellular C3 and C5 regulate vesicular trafficking of lysosomes^26^ and autophagosomes,^27^ respectively, and non-cytolytic MACs are associated with increased activity of heat-shock proteins^28^ and chaperones.^29^ These data posit a role for trafficking pathways in regulating non-cytolytic immunogenicity of MACs.

Alarmins are endogenous proteins that are basally non-immunogenic but become pro-inflammatory after losing their homeostatic compartmentalization^30^ or after becoming pathologically modified as is the case for amyloidogenic proteins.^31, 32^ We hypothesized that the burden of MAC proteins on ECs during CABMR could overwhelm endogenous mechanism(s) for protein homeostasis, facilitating formation of intracellular aggregates comprised of MAC-associated proteins. These aggregates could newly acquire intrinsic immunogenicity, thereby behaving similar to an alarmin. We interrogated our hypothesis using patient biospecimens, *in vitro* culture systems, and *in vivo* models. Our data provide a basis for understanding how MACs promote inflammation in the absence of widespread cell death.

## RESULTS

### MAC Aggregates Form Within Inflamed Patient Tissues

Complement component C9 has a propensity for polymerization. Within the MAC, ∼18 copies of C9 oligomerize to complete the pore.^33^ When resuspended in the absence of lipids and in the presence of Zn^2+^, C9 spontaneously polymerizes into tubular structures *in vitro*.^33, 34^ However, high concentrations of Zn^2+^ and low pH (4.5) inhibit tubular polymerization of C9, favoring nonspecific aggregation.^34^ In cell-free studies, we provisionally tested the propensity of C9 to form aggregates in comparison to other MAC-associated proteins. We found that, among MAC components, C9 showed the highest aggregate formation over time as labeled by thioflavin (Supplementary Figure 1*a*). C9 formed ring-like structures that increased in direct proportion to buffer acidity (Supplementary Figure 1*b*) and protein concentration (Supplementary Figure 1*c*).

Given the low pH environment and importance of the endolysosomal system in regulating intracellular Zn^2+^,^35^ we asked whether C9 may form aggregates within the endolysosomal system. To address this, we initially assessed for the presence of intracellular C9 aggregates in CABMR biopsies. To obviate confounding amyloid staining, we excluded *a priori* patients with primary amyloidosis as an indication for transplantation and post-transplant lymphoproliferative disorder (PTLD, Supplementary Table S1), a condition causing secondary amyloidosis.^3^ Immune-EM showed that adluminal cells in certain CABMR patients contained filamentous inclusions enriched for C9. These C9^+^ inclusions appeared to be contained within intracellular vesicles (Figure 1*a*). We did not detect filamentous inclusions in control tissues from transplant patients undergoing routine surveillance biopsies. In follow-up immunofluorescence (I.F). studies, we stained CABMR biopsies with Congo Red and thioflavin, clinically used dyes that quantify amyloid fibril formation^2^ but that also serve as a broad tool for detecting protein aggregation. Allograft vasculopathy (AV) is a vascular complication of CABMR, and vessels affected by AV show vascular laminations and adventitial scarring,^36, 37, 38^ sites known to contain elastin and collagen fibrils, respectively. We indeed noted strong thioflavin and Congo Red staining at these sites on AV vessels in 12 of 12 CABMR biopsies examined (Supplementary Figure 1*d*), precluding AV vessels as sites for analyzing the concurrent presence of protein aggregates.

**Figure 1.**
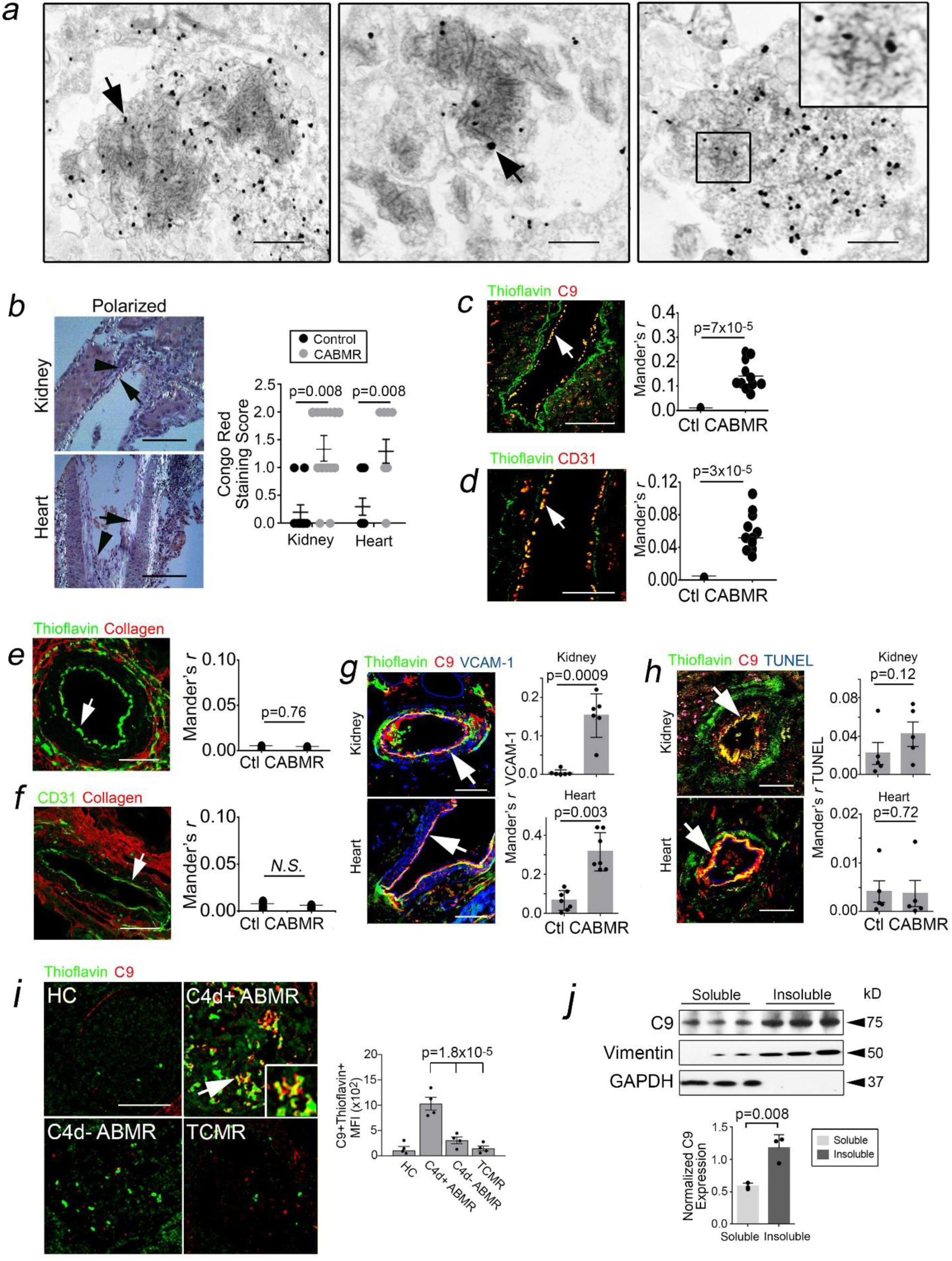
MAC Aggregates Form Within Inflamed Patient Tissues. C9 immune-EM of adluminal cells in CABMR biopsies (*a*). Congo Red staining was quantified in 4-8 high-power fields (hpfs) in kidney and heart CABMR biopsies. (*b*, n=12). Kidney and heart biopsies were analyzed by I.F. for thioflavin, C9, CD31, and collagen (*c-f,* n=12). Colocalization of VCAM-1 *(g,* n=6-7) and TUNEL (*h*, n=5) with C9^+^Thioflavin^+^ adluminal regions were quantified and averaged in 5 hpfs per sample. Healthy control (HC), C4d^+^ ABMR, TCMR, and C4d^-^ ABMR biopsies were analyzed by I.F (*i,* n=4 per group). Renal cell carcinoma (RCC) tissues were fractionated and analyzed for soluble and insoluble C9 normalized to GAPDH and vimentin, respectively (*j*, n=3). Data points indicate individual biopsy specimens analyzed (*b-j*). Scale Bar = 5μm (*a*), 200μm (*b-d, g-i*), and 100μm (*e,f*). Paired Student’s *t*-test (*b-h,j*) or two-way ANOVA followed by Tukey’s post-hoc correction (*i*).

However, in 5 of 12 CABMR biopsies we unexpectedly observed congophilic staining in adluminal cells within vessels lacking AV (Figure 1*b*). These adluminal cells co-stained for C9 and CD31 (arrows, Figure 1*c,d*) and moreover stained for thioflavin but not collagen (arrows, Figure 1*e,f*). C9^+^Thioflavin^+^ vessels expressed VCAM-1 (Figure 1*g*) at ∼3- to 10-fold higher levels than TUNEL staining (Figure 1*h*), indicating EC activation but not cell death. We secondarily surveyed C9^+^Thioflavin^+^ staining in glomeruli across various forms of ABMR in renal transplant recipients. Compared to ‘healthy control’ (HC) patients undergoing routine surveillance biopsies, C4d^+^ acute antibody-mediated rejection (ABMR), a form of rejection with strong complement activity, showed C9^+^Thioflavin^+^ glomerular staining (Figure 1*i*, arrow). In contrast, forms of rejection lacking strong complement activity, including C4d^-^ ABMR and T cell-mediated rejection (TCMR), showed comparatively lower C9^+^Thioflavin^+^ colocalization despite visualization of glomerular thioflavin staining. In further analyses, C4d^-^ ABMR tissues and TCMR showed strong thioflavin staining in vascular adventitia (Supplementary Figure 1*e*) and renal interstitium (Supplementary Figure 1*f,g*), likely reflecting regions of interstitial fibrosis.

To further test whether MAC-bound tissues contained C9 aggregates, we separately analyzed fresh tissues from patients with renal cell carcinoma (RCC, n=3). MAC deposition strongly occurred in RCC tissues relative to healthy controls (Supplementary Figure 1*h*). We fractionated RCC tissues heavily containing MACs into soluble and insoluble fractions, and we observed increased C9 within the SDS-insoluble pellet *vs* soluble supernatants (Figure 1*j*). Immune-EM, I.F., and tissue fractionation studies indicated that protein aggregates containing C9 may form in human tissues during inflammatory conditions involving chronic complement activity.

### The C9 Component of MACs Forms Intracellular Aggregates

To further model C9 aggregates, we used sera from allo-sensitized transplant candidates containing high titers of donor specific alloantibodies (DSA), termed patient reactive antibody (PRA).^24^ We previously found that PRA formed non-cytolytic MACs on HUVECs.^24^ As previously observed, PRA-treated HUVECs did not show increased cell death (Figure 2*a*). In immune-EM studies, PRA treatment for 6 hrs caused C9 to colocalize within large, perinuclear structures showing filamentous inclusions (Figure 2*b*), similar to patient CABMR biopsies. PRA caused thioflavin fluorescence to increase in both a dose-(left, Figure 2*c*) and time-dependent (right, Figure 2*c*) manner. In I.F. analyses, thioflavin showed punctate staining colocalizing with C9 in large, perinuclear vesicles (Figure 2*d*) lacking concurrent collagen fluorescence, thereby excluding confounding collagen staining as a cause for the observed thioflavin signals. We separated PRA-treated HUVECs into soluble and insoluble fractions and detected increased C9 within the SDS-insoluble pellet (Figure 2*e*) that, unlike pools of C9 within SDS-soluble supernatants, became resistant to mild proteinase K (PK) digestion (Supplementary Figure 2*a*). These results recapitulated ultrastructural, histologic, and biochemical findings observed in patient tissues.

**Figure 2.**
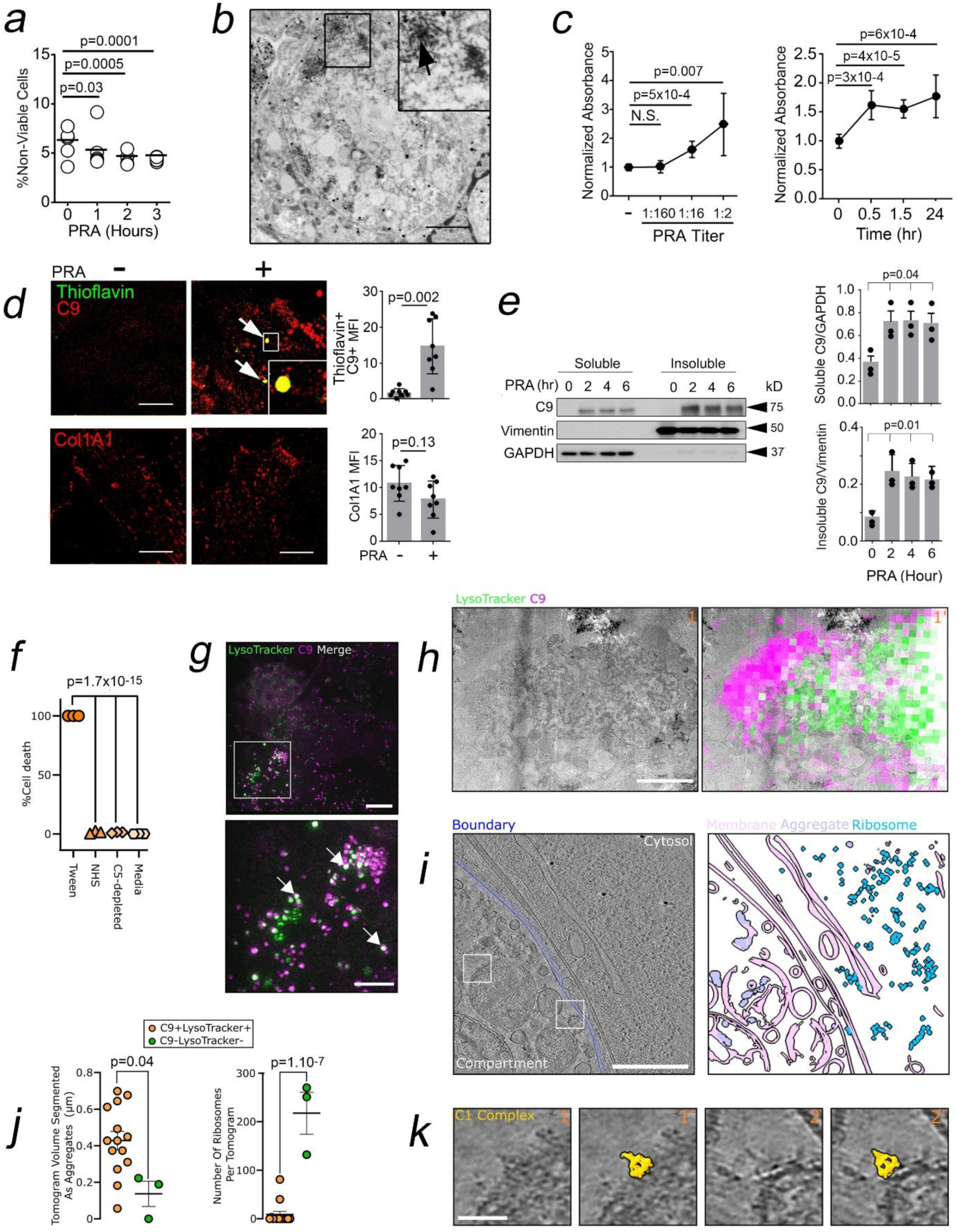
The C9 Component of MACs Forms Intracellular Aggregates. HUVECs pre-treated with IFN-γ (50ng/mL) for 48 hours were treated with PRA for various times prior to viability dye staining and analysis by FACS (*a*). Immune-EM for C9 (arrows) in HUVECs following PRA treatment for 6 hrs (*b*). IFN-γ-pretreated HUVECs were treated with PRA at various dilutions at 2 hrs (*c,* left) or at various times at 1:4 dilution (*c,* right) prior to assessing thioflavin fluorescence. By I.F., thioflavin but not collagen fluorescence was increased by PRA at 2 hrs (*d*). HUVECs were treated with PRA for various times, and soluble and insoluble fractions were analyzed by Western blot (*e*). HAP1 cells were incubated with either 1% Tween, media, 10% normal human serum (NHS), or 10% C5-depleted normal human serum, and cell death was assessed using Sytox (*f*). HAP1 cells treated with 10% C9-depleted serum, fluorescent C9 (purple), and LysoTracker (green) were plunge frozen and imaged using 3D cryoSIM. Representative max-intensity projection of cryoSIM z-stack is shown (*g,* top). Cropped region (white square) is enlarged to highlight C9^+^LysoTracker^+^ punctae (arrows, *g,* bottom). Representative 2D cryoCLEM images of milled lamellae from complement-treated HAP1 cells. The cryoEM image shows electron-dense aggregates and vesicles compartmentalized from the cytosol (*h*, left). The correlated fluorescence overlay shows signal for both C9 (purple) and LysoTracker (green) within the compartment (*h*, right). Images 1 and 1′ show the same field of view without and with the fluorescence overlay, respectively (*h*). Fluorescence and electron microscopy modalities were aligned using three-point reference alignment in the MAPS software. A 10 nm-thick slice from a cryo-electron tomogram collected from a C9^+^LysoTracker^+^ lamellae region (*i*, left). The lipid membrane boundary separating the aggregate-containing compartment from the cytosol is highlighted (blue). Automated segmentation of cellular features (membranes, pink; ribosomes, cyan; aggregates, purple) for the tomogram slice are displayed (*i*, right). Quantification of segmented features for all tomograms collected, including C9^+^LysoTracker^+^ (orange circles) and C9^-^LysoTracker^-^ (green circles) regions of the grid (*j*). Enlarged views of the regions highlighted by white boxes in panel *i*, showing densities consistent with the size and shape of previously reported C1 complexes (EMD-4232; yellow surface representation, *k*). Images 1 and 1′, and 2 and 2′, show the same fields of view without and with the fitted C1 density (EMD-4232), respectively (*k*). Data points indicate technical replicates for number of sample wells (*a,c,f*), number of cells (*d,*), or number of tomogram regions (*j*). Experiments repeated 3 times using different HUVEC donors (*d,e*). Error bars indicate standard deviations. Scale Bar =15μm (*b*), 50μm (*d*), 10μm (*g*, top), 5μm (*g,* bottom), 2μm (*h*), 0.1μm (*i*), 0.01μm (*k*). Paired Student’s *t*-test (*d*), one-way ANOVA with Tukey’s post-hoc correction (*a,c,e,f*), Welch’s *t*-test (left, *j*), and Mann-Whitney U-Test (right, *j*).

To identify culprit mediator(s) causing thioflavin fluorescence, we separated PRA sera into its IgG^+^ and IgG^-^ fractions and found that while these fractions separately showed minimal effects, combining the IgG^-^ and IgG^+^ fractions strongly potentiated thioflavin fluorescence (Supplementary Figure 2*b*). This ruled out IgG or contaminant(s) as a principal cause for increased thioflavin staining and suggested a role for complement activity. We next treated HUVECs with the IgG^+^ fraction of PRA combined with C9-deficient reference sera. IgG-induced complement activation with C9-deficient sera minimally increased thioflavin fluorescence compared to the IgG^+^ fraction alone (Supplementary Figure 2*c*, lane 3 *vs* lane 4). Addition of C9, which alone showed no effects (lane 1), significantly rescued thioflavin staining (lane 4 *vs* lane 5), indicating a role for C9 aggregates as a cause for thioflavin fluorescence.

To directly visualize internalized C9 in unfixed cells, we used focused ion beam and scanning electron cryo-microscopy (cryoFIB-SEM) together with cryo-electron tomography. We used a human cell line (HAP1 cells) which lacked two upstream complement regulators (CD46/CD55)^39^ to enhance terminal pathway activation and non-cytolytic conditions with human serum (Figure 2*f*). Cells were grown on cryo-EM grids and treated with C9-depleted serum supplemented with fluorescently labelled C9 and LysoTracker, a fluorescent marker used to label low pH (4,5) cellular compartments. Super resolution 3D structured illumination microscopy (cryo-SIM) of cells frozen on cryo-EM grids showed that internalized C9 co-localized with LysoTracker^+^ intracellular compartments (Figure 2*g*). Following cryoFIB milling, we investigated the nature of these compartments using correlative light and electron microscopy (cryo-CLEM, Figure 2*h*, Supplementary Figure 2*d*). In agreement with our patient-derived samples and PRA-treated HUVECs, we observed large, membrane-enclosed compartments comprised of dense, proteinaceous aggregates and smaller clustered vesicles. Next, fluorescence was used to target the collection of tomographic tilt series of areas that were C9^+^LysoTracker^+^. Control tilt series were collected from areas on the grid that were C9^-^LysoTracker^-^. We next performed automated cell feature segmentation (Figure 2*i*) which showed increased volume of aggregates within C9^+^LysoTracker^+^ tomograms (left, Figure 2*j*) and decreased ribosomes (right, Figure 2*j*), suggesting that aggregate-containing compartments are partitioned from the cytosol. Intriguingly, we observed densities consistent with C1 complexes on the surface of some vesicles within the membrane-bound compartment (Figure 2*k*). We did not observe densities consistent with MAC pores, suggesting that C9 aggregation is not related to the pore forming activity of C9. Filamentous aggregates were notably lacking in cells for which complement was basally activated. Therefore, our data suggested that while complement activation on nucleated cells trigger accumulation of C9 aggregates in endolysomal compartments, filamentous aggregates may be specific to the high protein burden caused by enhanced classical pathway activation on ECs, either during chronic complement activation (CABMR, RCC) or with PRA sera containing high titers of DSA. Together, these data supported the notion that complement activation on nucleated cells generates non-cytolytic aggregates of C9.

### NUMBL is a Rab35 Effector Promoting Intracellular C9

Based on EM findings, we asked whether C9 aggregates formed at the cell surface and/or intracellularly. We previously found that C9 became internalized via clathrin-mediated endocytosis (CME).^15^ Exploiting this, we pre-treated HUVECs with CME inhibitors, Dynasore and PitStop2, and tested effects on C9 aggregates. Both CME inhibitors visibly increased C9 levels at the cell surface (Supplementary Figure 2*e*, arrows), and this significantly reduced total thioflavin fluorescence in PRA-treated HUVECs. Pitstop2 significantly decreased insoluble C9:soluble C9 ratios (lane 2 *vs* lane 4, Supplementary Figure 2*f*), and these effects were phenocopied by siRNA *vs* dynamin-2 (DNM2), the EC-specific dynamin isoform causing membrane scission and endosome formation (Supplementary Figure 2*g*). We next stained PRA-treated HUVECs with a non-permeant dye staining protein aggregates (Proteostat) with or without cell permeabilization. Our data showed that incorporating the Triton-X permeabilization step potentiated Proteostat^+^C9^+^ staining by ∼200% (Figure 3*a*). These data together with cryo-CLEM studies (Figure *2g-j*), indicated that C9 aggregates formed intracellularly but not at the cell surface, posting a requirement for C9 internalization.

**Figure 3.**
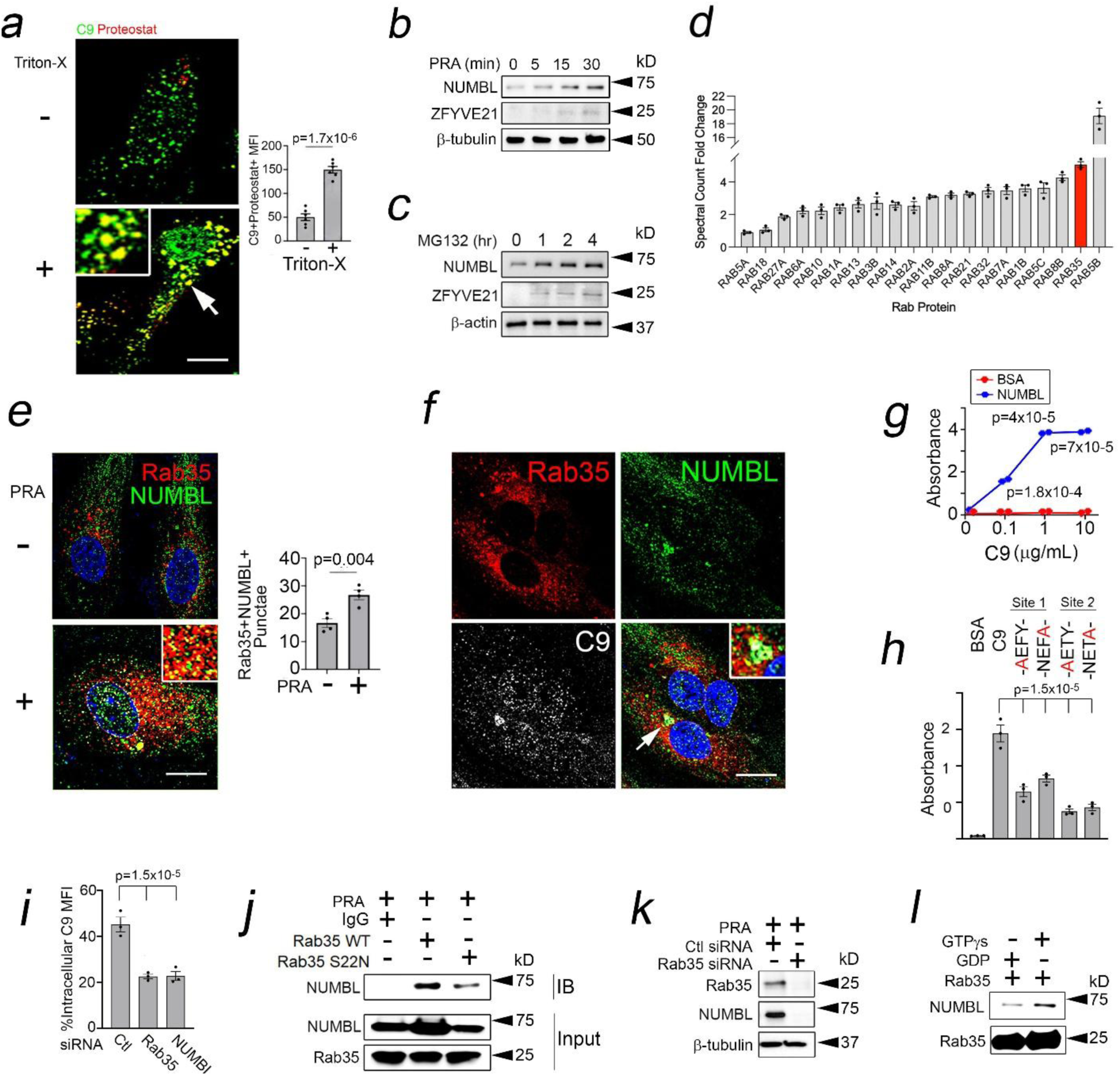
NUMBL is a Rab35 Effector Promoting Intracellular C9. HUVECs were treated with PRA for 1 hour, and I.F. was performed with or without a Triton-X permeabilization prior to I.F. staining (*a*). HUVECs were treated with PRA (*b*) or MG132 (*c*) and analyzed by Western blot. LC-MS/MS analysis of ZFYVE21-GFP vesicles (*d*). PRA-treated HUVECs were analyzed by I.F. (*e,f*). NUMBL protein was tested for binding to C9 (*g*) and C9 mutant proteins (*h*, 1μg/mL) by ELISA. HUVECs were treated with PRA for 1 hour, and intracellular C9 was assessed by FACS with siRNA against Rab35 or NUMBL (*i*). Pulldowns of Rab35 (*j*) and Western blots of PRA-treated HUVECs (*k*). GTPγs or GDP was incubated with Rab35 for 30 min prior to addition of NUMBL for 30 min at RT prior to Western blot (*l*). Data points indicate technical replicates for number of cells analyzed (*a,e*) or number of sample wells (*g-i*). Error bars indicate standard deviations. Scale Bar = 30μm (*a,e-f*). Experiments repeated n=3 (*b,c,j-l*) times. Paired Student’s *t*-test (*a,e*) or one-way ANOVA with Tukey’s post-hoc correction (*g-i*).

Based on the above, we embarked on studies to identify mediator(s) of C9 internalization. Early endosomes are enriched in adaptor proteins that bind surface cargoes like C9 to mediate their internalization. To uncover such protein(s), we performed LC-MS/MS of ZFYVE21^+^ endosomes isolated from PRA-treated HUVECs. ZFYVE21 is a highly conserved protein that we chose as a target for LC-MS/MS analyses based on our prior study showing its high colocalization with early endosomes,^22^ sites where C9 may become internalized. Across 3 runs, LC-MS/MS yielded 3,347 unique proteins, 586 of which were uniquely upregulated by PRA at the 30 min timepoint (Supplementary Figure 3*a*). These proteins included MAC proteins (C5b/6/7/8α/8γ/9) and inflammasome proteins (NLRP3, casp1) identified in our prior proteomic datasets.^22, 23^ Among upregulated proteins, we identified endocytic adaptors whose consensus binding motifs appeared on C9. From this, we selected molecules that strongly modulated NF-κB in a prior genome-wide siRNA screen of PRA-treated HUVECs.^15^ This iterative search identified NUMBL, an endocytic adaptor whose binding motif [F-X-N-P/X-X-Y or N-P/X-X-Y, X=any amino acid]^40^ appeared at 2 sites on C9 (Supplementary Figure 3*b*).

NUMBL is a conserved and pleiotropic protein with functional roles in both endocytosis and activation of NF-κB.^41, 42, 43^ To begin to define its roles in C9 internalization, we initially examined NUMBL expression in PRA-treated HUVECs. With PRA, NUMBL became upregulated within <15 minutes (Figure 3*b*). We treated HUVECs with MG132 and found that NUMBL protein became potentiated over time, similar to ZFYVE21,^22, 23^ indicating that NUMBL levels are post-translationally regulated via proteasome degradation (Figure 3*c*). Following its post-translational stabilization after PRA treatment, we addressed site(s) where NUMBL colocalized. Rab35^+^ endosomes canonically form upstream of early endosomes and, like NUMBL, Rab35+ endosomes show functional roles in both endocytosis^44^ and inflammation,^45^ positing relationships with NUMBL. Among Rab proteins, Rab35 was strongly enriched within ZFYVE21^+^ vesicles in LC-MS/MS (Figure 3*d*), and among endosome populations tested, NUMBL most heavily colocalized with Rab35^+^ endosomes at the 15 min timepoint (Figure 3*e*), forming Rab35^+^NUMBL^+^ punctae colocalizing with C9 (Figure 3*f*).

Based on I.F. data, we tested NUMBL interactions with C9, focusing on NUMBL’s canonical binding motifs. In ELISAs, NUMBL but not bovine serum albumin (BSA) showed dose-dependent binding to C9 (Figure 3*g*), and NUMBL binding was significantly reduced following alanine substitutions affecting either of NUMBL’s canonical binding motifs on C9 (Figure 3*h*). By FACS, intracellular C9 was reduced by ∼50% with NUMBL siRNA (Figure 3*i*), together indicating that NUMBL directly binds C9 to mediate C9 internalization.

NUMBL is post-translationally stabilized by PRA (Figure 3*b,c*) concurrent with its recruitment to Rab35^+^ endosomes (Figure 3*e*). Based on this, we examined secondary interactions between NUMBL and Rab35. Rab35 dominant negative (DN) carries an S22N mutation, locking Rab35 in its inactive GDP-bound form. We pulled down Rab35 endosomes in PRA-treated HUVECs and found that Rab35 DN blocked recruitment of NUMBL to Rab35 endosomes (lane 2 *vs* lane 3, IB, Figure 3*j*), and this reduced total cellular levels of NUMBL (lane 2 *vs* lane 3, Input, Figure 3*j*). With PRA, siRNA-mediated loss of Rab35 similarly reduced NUMBL levels (Figure 3*k*). Rab5 activation occurs downstream of Rab35,^44^ and we found that Rab5 DN (S23N), while strongly reducing levels of ZFYVE21, a Rab5 effector,^22^ showed weaker effects on levels of NUMBL (lane 2 *vs* lane 4, Supplementary Figure 3*c*), indicating a selective, Rab35-mediated response. These data showed that Rab35^+^ endosomes sequestered NUMBL to post-translationally increase NUMBL protein levels.

Rab proteins complexed to GTP selectively bind effector molecules to execute downstream function(s). Given the ability of Rab35 activity to modulate total cellular levels of NUMBL (Figure *3j,k*), we tested whether NUMBL may bind Rab35-GTP as an effector molecule. We incubated recombinant Rab35 with non-hydrolyzable GTPγS or GDP prior to addition of recombinant NUMBL and found that NUMBL showed selective binding to active Rab35-GTP complexes (Figure 3*l*). Based on its selective binding to Rab35-GTP and its functional role in mediating C9 internalization, our data operationally defined NUMBL as a Rab35 effector protein. In sum, our data showed that Rab35^+^ endosomes capture NUMBL, shielding this same protein from proteasome degradation and rapidly enhancing its stability. Upon its post-translational stabilization, NUMBL functions as a Rab35 effector, heavily colocalizing with Rab35+ endosomes where it binds C9 to promote C9 internalization.

### NUMBL Promotes C9 Aggregates within Rab7^+^ Endosomes

Extending our cryo-CLEM data in Figure 2, we analyzed the fate of internalized C9 within endolysosomes whose acidified microenvironments we posit favored C9 aggregation. We initially used live cell imaging to visualize entry of extracellular C9 into acidified endosomes. We stably transduced HUVECs with a Rab5-GFP reporter and treated these cells with PRA spiked with C9 labeled with pHrodo, an acidic pH-sensitive dye. Compatible with our cryoSIM data (Figure 2*g*), we found that pHrodo^+^Rab5^+^ punctae dramatically increased within 50 min, indicating that upon its internalization C9 became exposed to acidic pH, a condition known to induce nonspecific aggregation of C9 (Figure 4*a*, Supplementary Movie 1).^34^ To identify intracellular compartment(s) where C9 aggregation occurred, we performed pulse-chase studies and detected C9^+^Thioflavin^+^ punctae within early endosomes (Rab5), late endosomes (Rab7), and lysosomes (LAMP1, Figure 4*b*). Among these populations, Rab7^+^ endosomes showed highest staining for both C9 and thioflavin while LAMP1^+^ lysosomes showed lower staining. We consolidated the above using vesicular flow cytometry via our published protocol.^22, 23^ After gating on C9^+^ vesicles, we noted basal thioflavin staining in ∼10-15% of vesicles harvested from untreated cells. Basal thioflavin staining predominantly occurred in Rab7+ compartments, possibly a reflection of fibrillar components and/or substrates undergoing basal turnover. With PRA, thioflavin staining increased across all endolysosomal populations examined and was ∼40-45% greater in Rab7^+^ vesicles compared to Rab5^+^ and LAMP-1^+^ vesicles (Figure 4*c*). Our data show that C9 aggregates form within the endolysosomal pathway, particularly within Rab7^+^ endosomes.

**Figure 4.**
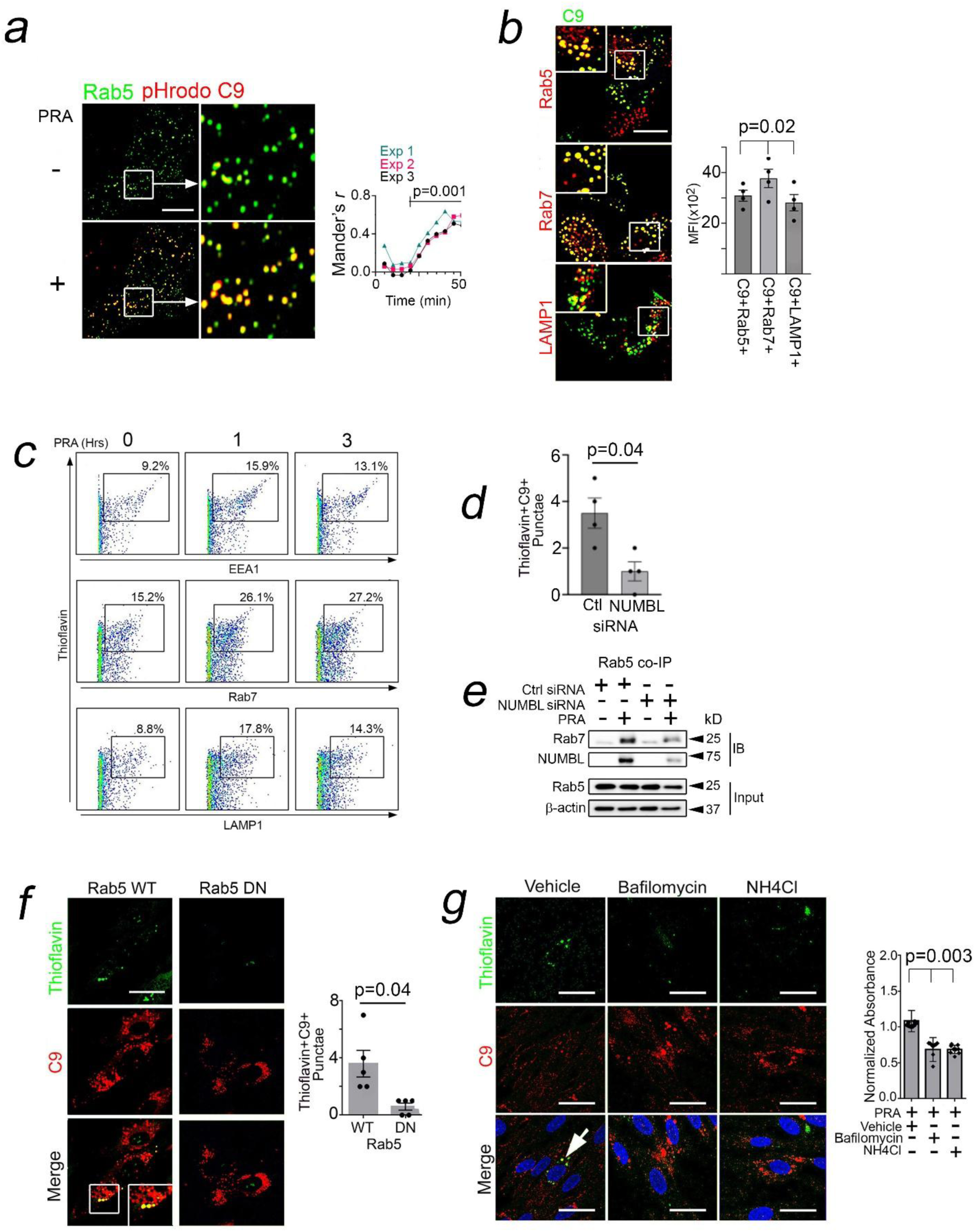
NUMBL Promotes C9 Aggregates within Rab7^+^ Endosomes. Rab5-GFP HUVECs were treated with PRA spiked with pHrodo-labeled C9 and analyzed by live cell imaging (*a*). HUVECs were treated with PRA for 1 hour, washed, and analyzed 1 hour afterwards by I.F. (*b*). PRA-treated HUVECs were sonicated, and vesicular fractions were enriched by sucrose gradient centrifugation then analyzed by FACS at the times indicated (*c*). PRA-treated HUVECs were analyzed by I.F. (*d*) or by Western blot following Rab5 pulldowns at 1 hour (*e*). HUVECs were transduced with Rab5 WT and Rab5 DN constructs, treated with PRA for 2 hrs prior to I.F., and punctae per cell were quantified (*f*). HUVECs were exposed to vehicle, bafilomycin (100 nM), or NH_4_Cl (10 μM) for 2 hours, PRA was added, and thioflavin fluorescence was assessed 4 hours later (*g*). Data points indicate technical replicates for cell regions (*b*) and cell punctae (*d,f,g*). Experiments repeated n=2 (*e*), n=3 (*a,c,d,f,g*), or n=5 (*b*) times using different HUVEC donors. Error bars indicate standard deviations. Scale Bar = 30μm (*a,b*), 20μm (*f,g*). One-way ANOVA with Tukey’s post-hoc correction (*a,b,g*) or paired Student’s *t*-test (*d,f*).

To test roles for NUMBL in mediating C9 aggregates, we performed NUMBL knockdowns and noted that NUMBL siRNA significantly reduced C9^+^Thioflavin^+^ punctae in PRA-treated HUVECs (Figure 4*d*). We pulled down Rab5+ endosomes using co-immunoprecipitations (co-IPs) and found that NUMBL siRNA strongly reduced the population of transitioning endosomes acquiring Rab7 (Figure 4*e*). Rab5 activity is required for acquisition of Rab7, and based on the above, we tested effector functions of Rab5 for mediating C9 aggregates. We found that blocking Rab5 activity with Rab DN (S23N) virtually abolished thioflavin staining in PRA-treated HUVECs (Figure 4*f*), and blocking vesicular acidification, a Rab5-dependent process, with NH4Cl buffer alkalinization or with bafilomycin, a V-ATPase inhibitor, significantly reduced thioflavin staining when compared to controls (Figure 4*g*). Together these data indicated that NUMBL promoted C9 aggregates by modulating Rab5 activity and that loss of vesicular acidification, a Rab5-dependent process, ablated C9 aggregates. These findings are consistent with C9’s propensity for forming nonspecific aggregation under acidic pH in cell-free studies.^34^

### C9 Aggrephagy Induces NF-κB Activity and Causes EC Activation

Aggrephagy is a form of selective macroautophagy enabling degradation of protein aggregates that might otherwise cause cytotoxic effects.^46^ In this process, aggrephagy may activate inflammatory signaling including NF-κB.^46, 47^ C9 and thioflavin signals were detected in LAMP1+ degradative lysosomes (Figure 4*b,c*) in association with EC activation (Figure 1*g*), prompting us to ask whether, as a process, aggrephagy allowed C9 aggregates to acquire pro-inflammatory properties. Analysis of cellular compartments within C9^+^LysoTracker^+^ tomograms revealed double membrane structures that morphologically resembled phagophores (arrows, left, Figure 5*a*)^48^ and that, upon segmentation, heavily contained morphologic aggregates (right, Figure 5*a*). On this basis, we examined C9 aggrephagy in PRA-treated HUVECs. With PRA, HUVECs showed increased autophagic flux, marked by LC3-II and P62 (Figure 5*b*). Exploiting the fact that GFP fluorescence quenches at pH <5.0, the ‘traffic light’ reporter (LC3B-GFP-RFP) generates GFP^+^RFP^+^ signals marking non-acidified autophagosomes, and GFP^-^RFP^+^ signals marking acidified autophagolysosomes. In ‘traffic light’ HUVECs, PRA caused increased GFP^+^RFP^+^ LC3B punctae, indicating increased generation of autophagosomes (Figure 5*c*), and in co-IPs, we observed that C9 became ubiquitinylated (Supplementary Figure 4*a*), the first step towards degradative targeting to autophagolysosomes. At ∼2 hrs, C9 became contained within LC3B^+^ membrane cages (Supplementary Figure 4*b*) displaying aggresome markers (Figure 5*d*, Supplementary Figure 4*c-e*). Blocking autophagosome-lysosome fusion with chloroquine (CQ) caused gross enlargement of C9^+^ vesicles (Supplementary Figure 4*f*), indicating that C9 had trafficked through a macroautophagic pathway. We performed pulse-chase studies in PRA-treated HUVECs and found that siRNA depletion of autophagy mediators, ATG5 (Figure 5*e*) or ATG16L (Supplementary Figure 4*g*), increased intracellular C9. We consolidated the above using CRISPR/Cas9-mediated knockouts of ATG16L in HEK293 cells. Phenocopying effects of siRNA, genetic loss of ATG16L ablated LC3-II while increasing cellular levels of C9 (Supplementary Figure 4*h*). These data together indicated that C9 aggregates become substrates for aggrephagy.

**Figure 5.**
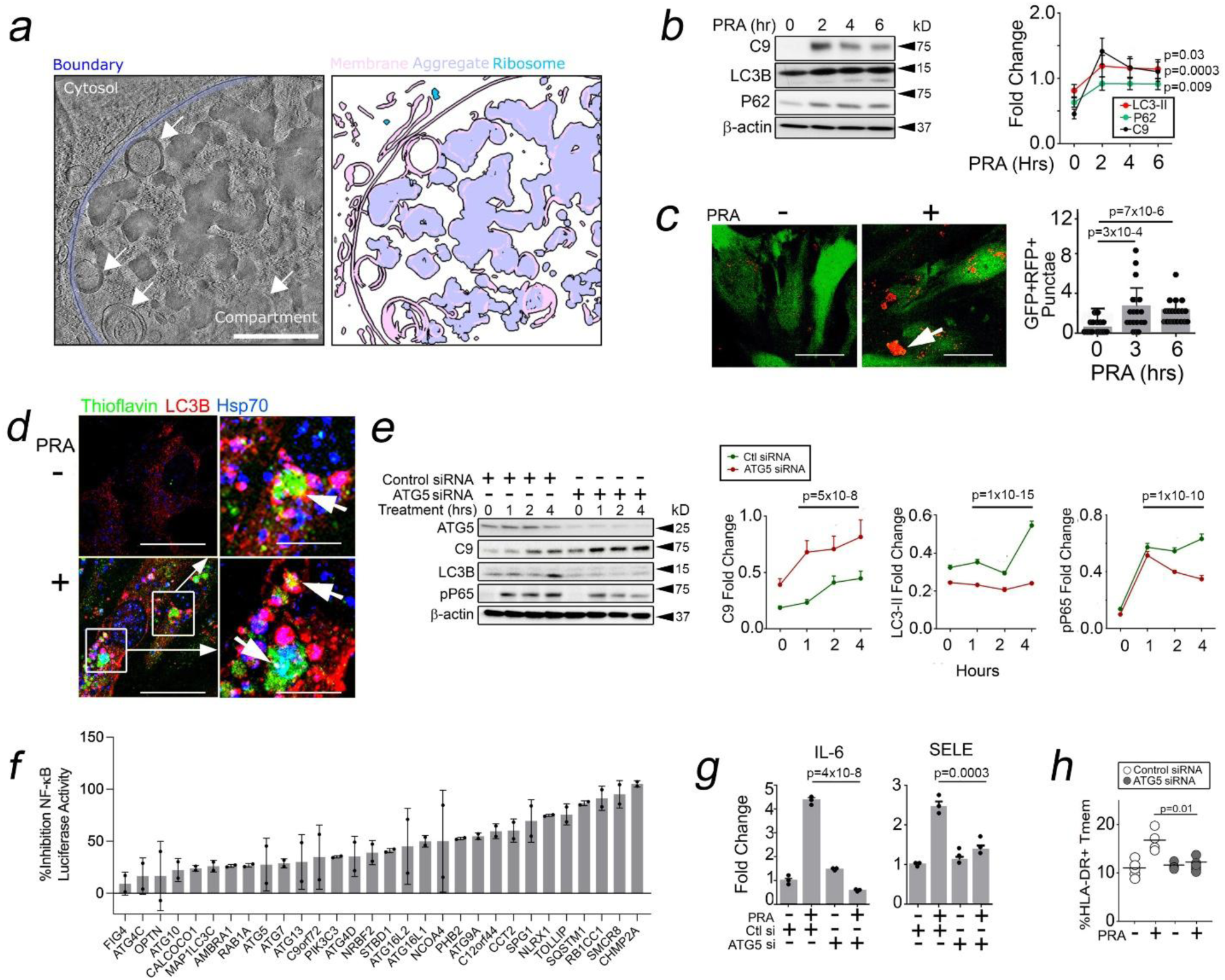
C9 Aggrephagy Induces NF-κB Activity and Causes EC Activation. HAP1 cells grown on cryoEM grids and treated with C9-depleted serum supplemented with fluorescent C9 and LysoTracker. Grids were imaged and analyzed as described in Figure 2. A 10 nm-thick slice from a cryo-electron tomogram collected from a C9^+^LysoTracker^+^ lamellae region (*a*, left). Double-membrane cellular structures morphologically consistent with phagophores (arrows) with the membrane boundary separating the aggregate-containing compartment from the cytosol highlighted (blue). Automated segmentation of cellular features for the tomogram slice are displayed (*a*, right). HUVECs were treated with PRA for the times indicated prior to Western blot analysis (*b*). HUVECs transduced with mRFP-GFP-LC3B were treated with PRA and analyzed for GFP^+^RFP^+^ punctae (*c*). HUVECs were pre-treated with chloroquine (CQ) for 30 minutes prior to PRA treatment for 2 hours (*d*). HUVECs were transfected with siRNA as indicated and analyzed in pulse-chase studies after PRA treatment (*e*). HUVECs were stably transduced with an NF-κB luciferase reporter,^15^ subjected to siRNA knockdown as indicated, and treated with PRA for 6 hours prior to assessing luciferase activity (*f*). HUVECs were transfected with siRNA against ATG5 and treated with PRA for 4 hours prior to performing qRT-PCR (*g*). HUVECs transfected with siRNA against ATG5 were cocultured for 10 days with alloimmune CD4^+^CD45RO^+^ T cells, and T cells were harvested for FACS analysis (*h*). Data points indicate cells analyzed (*c*), biological replicates (*b,e,g*), or technical replicates (*h*). Experiments repeated ≥3 times using different HUVEC donors. Error bars indicate standard deviations. Scale Bar = 0.1μm (*a*), 30μm (*c*) and 20μm (*d*) One-way ANOVA (*c,g-h*) or two-way ANOVA (*b,e*) with Tukey’s post-hoc correction.

To test whether aggrephagy conferred immunogenic properties to C9 aggregates, we transduced HUVECs with an NF-κB activity luciferase reporter and performed a limited siRNA screen against genes annotated to ‘selective macroautophagy’ (KEGG). This analysis showed that knockdown of 30 of 52 genes involved in macroautophagy significantly inhibited NF-κB activity in PRA-treated HUVECs (Figure 5*f*). A re-analysis of Western blots in Figure 5*e* and Supplementary Figure 4*g* further showed that siRNA against autophagy genes in PRA-treated HUVECs reduced phosphoP65 (pP65), and this significantly reduced NF-κB-dependent gene transcripts (Figure 5*g*, Supplementary Figure 4*i*).^24^ Human ECs express sufficient costimulatory molecules to directly prime CD4^+^CD45RO^+^ memory T cells (Tmem) in EC:T cell cocultures,^24^ and ECs transfected with siRNA *vs* autophagy genes showed significantly reduced ability to elicit activation (HLA-DR), effector (IFN-γ) and proliferative (CFSE dilution) responses in alloimmune Tmem (Figure 5*h*, Supplementary Figure 4*j*). These data supported the notion that, as a process, aggrephagy conferred C9 aggregates with pro-inflammatory effects.

### NUMBL Stabilizes ZFYVE21 to Promote C9 Aggrephagy

We sought to define a mechanism linking NUMBL to C9 aggrephagy. ZFYVE21 is a conserved, endosome-associated protein^22, 49^ that we previously found could function as a Rab5 effector modulating NF-κB.^22, 23^ Based on the ability of NUMBL to modulate Rab5 (Figure 4*e*), we asked whether NUMBL utilized ZFYVE21 to mediate C9 aggrephagy. ZFYVE21 became upregulated in PRA-treated HUVECs concurrent with LC3-II (Figure 6*a*), and ZFYVE21 levels were reduced by disrupting Rab35-NUMBL complexes, either with Rab35 DN (Figure 6*b*) or NUMBL siRNA (Figure 6*c*). These data identified ZFYVE21 as a potential downstream effector by which NUMBL might modulate aggrephagic flux.

**Figure 6.**
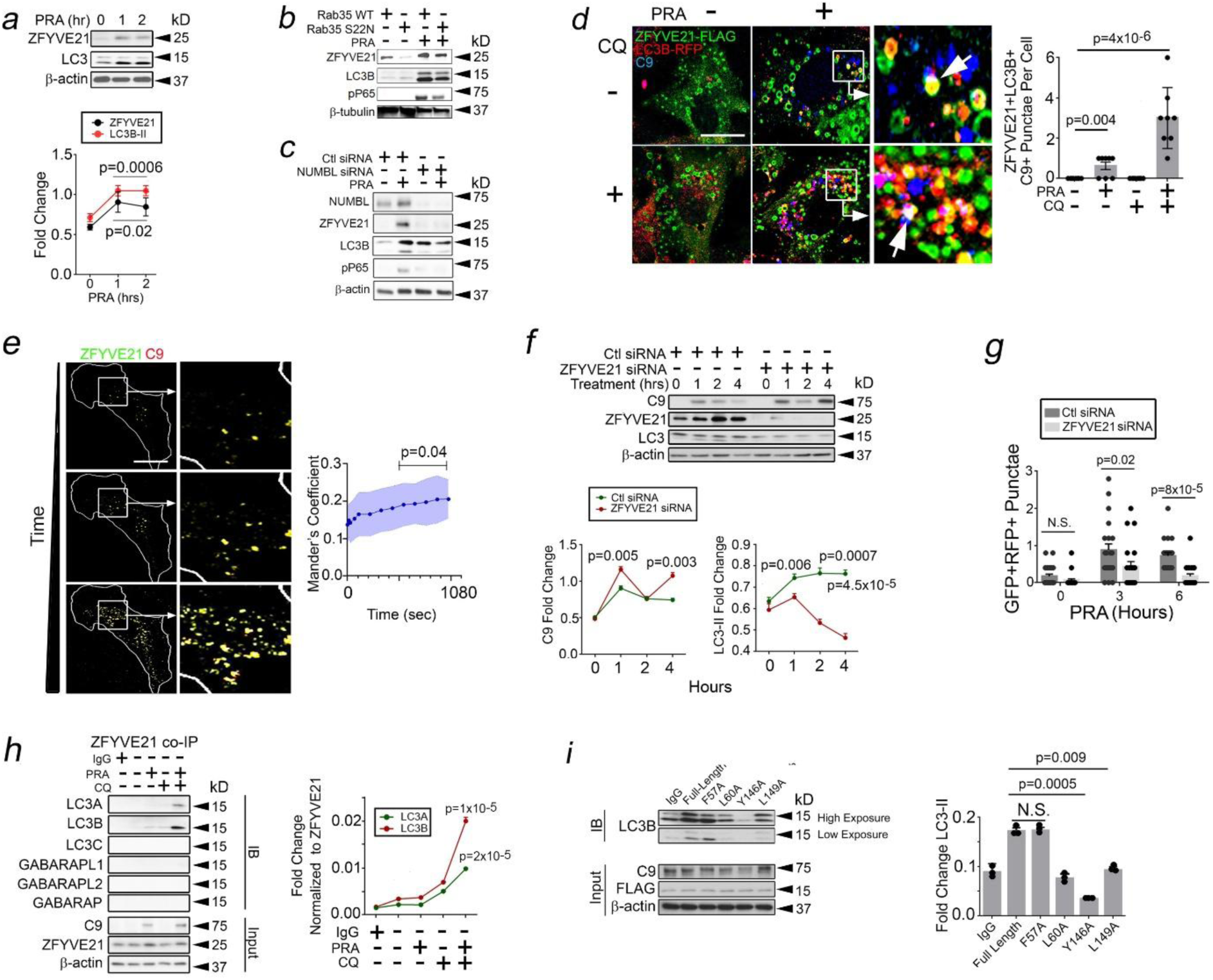
NUMBL Stabilizes ZFYVE21 to Promote C9 Aggrephagy. HUVECs were treated with PRA and analyzed by Western blot (*a-c*). HUVECs transduced with ZFYVE21-FLAG and LC3B-RFP were treated with PRA containing C9-AF647 protein (10μg/mL) and either with or without CQ for 2 hours prior to analysis by I.F. (*d*). HUVECs transduced with ZFYVE21-GFP were treated with PRA containing AF647-labeled C9 prior to live cell imaging (*e*). HUVECs were transfected with ZFYVE21 siRNA and analyzed in pulse-chase studies at various times after PRA treatment (*f*). HUVECs transduced with mRFP-GFP-LC3 constructs and transfected with ZFYVE21 siRNA were analyzed by I.F. after PRA treatment for the indicated times (*g*). HUVECs transduced with ZFYVE21 were treated with PRA and with or without CQ prior to ZFYVE21 co-IPs and Western blot analysis (*h*). HUVECs were transduced with FLAG-tagged ZFYVE21 constructs containing alanine substitutions at the amino acid sites indicated prior to FLAG co-IPs and Western blot analysis (*i*). Data points indicate technical replicates for cell punctae (*d,g*) or sample wells (*i*). Experiments were repeated n=2 (*b,c,h,i*) or n=3 (*a,d,e,f*) times using different HUVEC donors. Error bars indicate standard deviations. Scale Bar = 30μm (*d*) or 40μm (*e*). One-way ANOVA with Tukey’s post-hoc correction (*a,d,e,f,h,i*) or paired Student’s *t-*test (*g*).

To test this, we initially treated HUVECs with PRA spiked with AF647-labeled C9 protein, a treatment forming fluorescent C9^+^ vesicles.^22^ At the 1hr timepoint, ZFYVE21 heavily colocalized with Thioflavin^+^ aggregates (arrow, Supplementary Figure 5*a*). The number of C9^+^ZFYVE21^+^ punctae containing LC3B were grossly increased with CQ (Figure 6*d*), indicating that ZFYVE21 had trafficked through a macroautophagic pathway. To directly visualize this, we performed live cell imaging and observed increased C9^+^ZFYVE21^+^ vesicles over time in HUVECs treated with PRA spiked with AF647-labeled C9 (Figure 6*e*, Supplementary Movie 2). C9^+^ZFYVE21^+^ vesicles showed increased displacement and size (Supplementary Figure 5*b*), behaviors compatible with maturing vesicles prior to their entry into the aggrephagy pathway.^50^

In pulse-chase studies, ZFYVE21 siRNA ablated autophagic flux while potentiating C9 (Figure 6*f*), and this ablated GFP+RFP+ autophagosomes in ‘traffic light’ HUVECs treated with PRA (Figure 6*g*). Phenocopying ZFYVE21 siRNA, gene deletion of ZFYVE21 via CRISPR/Cas9 ablated LC3-II while potentiating C9 (Supplementary Figure 5*c*). We found that in contrast to PRA, serum starvation induced autophagic flux in ECs without upregulating ZFYVE21 (Supplementary Figure 5*d*), and genetic loss of ZFYVE21 showed no effects on starvation-induced LC3-II (lanes 2-4 *vs* lanes 6-8, Supplementary Figure 5*d*). These data indicated a role for ZFYVE21 in aggrephagy of C9, a form of selective macroautophagy, but not starvation-induced, non-selective macroautophagy.

To address a mechanism by which ZFYVE21 mediated aggrephagy, we tested whether ZFYVE21 could interact with one or more ATG8 family proteins. ATG8 family proteins critically anchor signaling proteins to aggresomes and are critically required for generating autophagosome membranes.^51^ We performed pulldowns in ZFYVE21-FLAG ECs treated with CQ to block degradation of ATG8 family proteins which might otherwise become substrates for aggrephagy.^52, 53^ In co-IPs, ZFYVE21 principally interacted with LC3A and LC3B (Figure 6*h*). LC3A/B are ATG8 family proteins that bind adapter proteins via LC3 Interacting Regions (LIRs, W/F/Y-X-X-I/L/V, X = any amino acid) and recently identified GABARAP interacting motifs (GIMs, W/F-V/I-X-V).^53^ While ZFYVE21 did not contain GIMs, we identified 2 LIRs between AA57-60 and AA146-149 (Supplementary Figure 5*e*). Alanine substitutions of ZFYVE21 at these 2 LIRs showed that L60, Y146, and L149 were required for ZFYVE21 binding to LC3B (Figure 6*i*). Together, these data identify ZFYVE21 as an LC3B interacting protein utilized by NUMBL to promote C9 aggrephagy.

### ZFYVE21-RNF34-P62 Complexes Mediate C9 Aggrephagy

With PRA, C9 became ubiquitinylated (Supplementary Figure 4*a*) and targeted for aggrephagy (Figure 5*e*, Supplementary Figure 4*g*). Based on this, we sought to identify E3 ubiqutin ligase(s) utilized by ZFYVE21 to mediate C9 aggrephagy. To do this, we conducted a second iterative search of LC-MS/MS data obtained from ZFYVE21^+^ vesicles from Figure 3. We searched for proteins upregulated by PRA containing RING and HECT domains, domains known to show E3 ubiquitin ligase activity, and we cross-referenced this list against proteins showing *z*-scores ≥2 in a prior genome-wide siRNA screen for NF-κB.^15^ Among proteins identified, we focused on RNF34 as this protein contains both a FYVE domain allowing colocalization to endosomes and a RING domain enabling ubiquitinylation.^54, 55^

As RNF34 has not been previously connected to autophagic processes, we first considered whether RNF34 interacted with ZFYVE21 within aggresomes. RNF34 pulled down with ZFYVE21 along with C9 and P62, an autophagy mediator (Figure 7*a*). At the 30 min timepoint, we previously found that RNF34 interacted with ZFYVE21 to modulate activity of inflammasomes.^23^ We found that RNF34 interactions persisted up to 2-4 hrs, a time when aggresome formation occurred. At these times, RNF34 heavily colocalized within ZFYVE21^+^Thioflavin^+^ aggresomes (Figure 7*b*). RNF34 siRNA reduced C9 ubiquitinylation while potentiating levels of C9 (Figure 7*c*), and this had the effect of attenuating the intrinsic immunogenicity of C9 for activating NF-κB (Figure 7*c*) and for promoting EC-mediated T cell activation (Figure 7*d*). Based on this, we tested whether RNF34 could ubiquitinylate C9. Among E2 conjugating enzymes tested, we selected UBCH5a due to its strong activity (Supplementary Figure 6*a*). We found that UBCH5a allowed RNF34 to directly K48 ubiquitinylate C9 in a manner requiring its RING domain containing E3 ubiquitin ligase activity (Figure 7*e*). We subsequently co-transfected CQ-treated HUVECs with RNF34 siRNA in the presence of either Ub-WT or Ub-DN constructs and found that RNF34 siRNA, like Ub-DN, blocked C9 ubiquitinylation (Figure 7*f*). These data indicated that RNF34 directly ubiquitinylates C9 to promote C9 aggrephagy.

**Figure 7.**
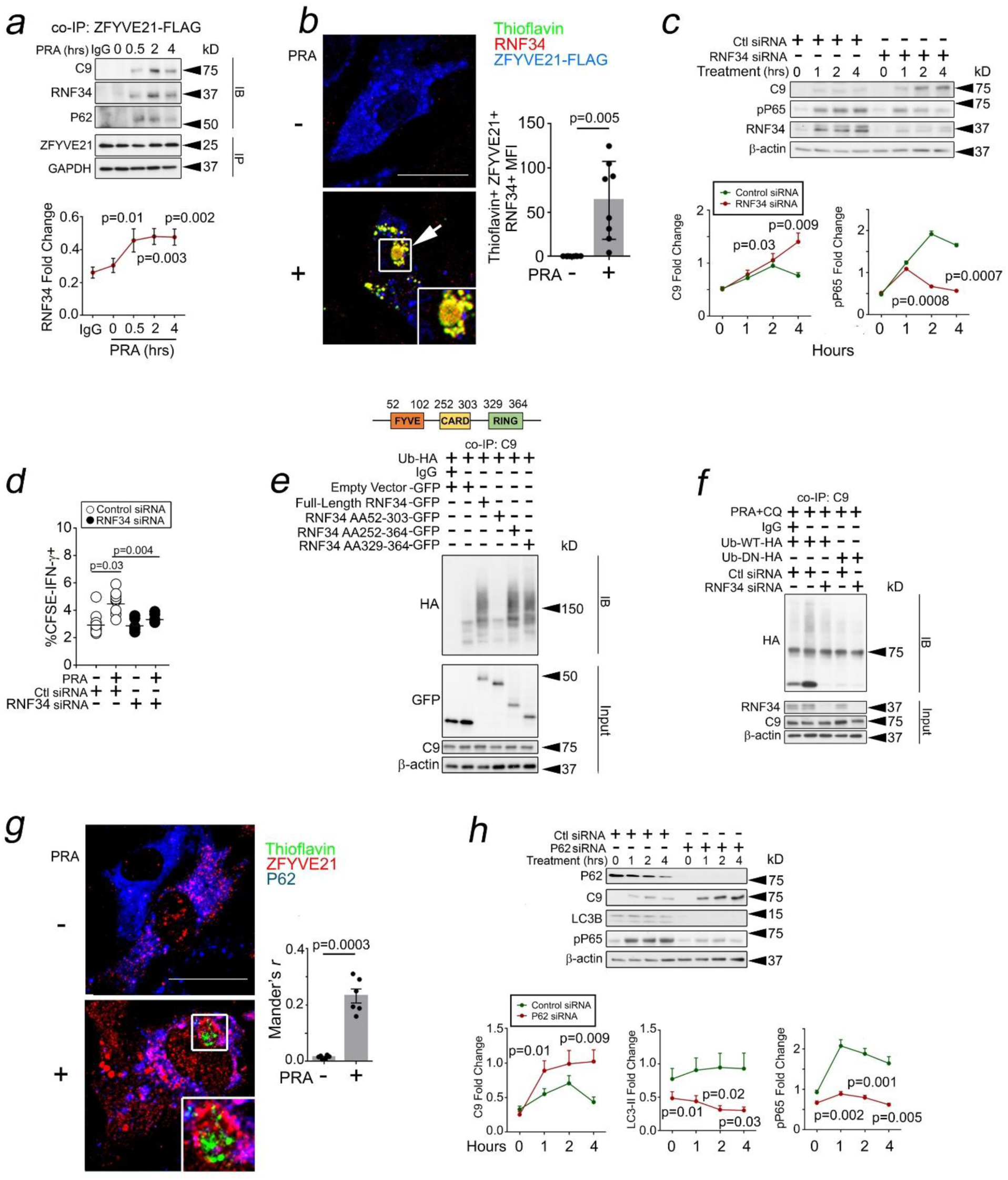
ZFYVE21-RNF34-P62 Complexes Mediate C9 Aggrephagy. HUVECs transduced with ZFYVE21-FLAG were treated with PRA at the indicated times prior to FLAG co-IP and Western blot analysis (*a*). HUVECs transduced with RNF34-HA were treated with PRA for 2 hours prior to staining for thioflavin and ZFYVE21 (*b*). HUVECs transfected with control siRNA or RN34 siRNA were analyzed in pulse-chase studies after PRA treatment (*c*). HUVECs transfected with siRNA against RNF34 were treated with or without PRA for 4hrs prior to coculture with alloimmune CD4^+^CD45RO^+^ T cells. T cells were harvested and analyzed by FACS 10-14 days later (*d*). HUVECs were transfected with Ub-HA and GFP-tagged constructs encoding various regions of RNF34 prior to Ub-HA co-IPs (*e*). HUVECs were co-transfected with Ub-WT and Ub-DN along with control or RNF34 siRNA, treated with PRA, and analyzed by Western blot following C9 co-IP (*f*). HUVECs were treated for PRA for 2hrs prior to I.F. analysis (*g*). HUVECs were transfected with P62 siRNA and analyzed in pulse-chase studies following PRA treatment (*h*). Data points indicate technical replicates for cell regions (*b,g*) or sample wells (*d*). Experiments repeated n=2 (*e,f*) or n=3 (*a,c,h*) times using different HUVEC donors. Error bars indicate standard deviations. Scale Bar = 20μm (*b,g*). One-way ANOVA (*a,d*) or two-way ANOVA (*c,h*) with Tukey’s post-hoc correction or paired Student’s *t*-test (*b,g*).

For aggrephagy to occur, ubiquitinylated substrates require binding to proteins containing ubiquitin binding domains (UBDs).^52^ ZFYVE21 and RNF34 lack UBDs, and in cell-free studies neither of these proteins showed binding to K48 Ub chains (Supplementary Figure 6*b*). In contrast, P62 showed robust binding to ubiquitin (Supplementary Figure 6*b*) and was found to pull down with ZFYVE21 (Figure 7*a*) and to colocalize with ZFYVE21^+^Thioflavin^+^ punctae (Figure 7*g*). In high exposure blots, our pulse-chase protocol caused P62 to decline over time in control cells, compatible with prior observations that P62 itself becomes an autophagic substrate (Figure 7*h*).^52^ P62 siRNA potentiated C9 while inhibiting LC3-II (Figure 7*h*), and this significantly decreased phosphoP65 in PRA-treated HUVECs (Figure 7*h*) as well as NF-κB luciferase activity (SQSTM1, Figure 5*f*). Collectively, our data support a sequence where (1) C9 aggregates are ubiquitinylated by RNF34 (Figure 7*e,f*), (2) ubiquitinylated C9 binds P62 (Figure 7*a*, Supplementary Figure 6*b*), and (3) C9-RNF34-P62 complexes are targeted to LC3B+ aggresomes by ZFYVE21 (Figure 6*f,g*, Figure 7*a-b,g*). In this sequence, ZFYVE21, an LC3B-interacting protein, appears to function as an adaptor, bridging C9-RNF34-P62 complexes to LC3B^+^ aggresomes.

### Signaling Responses Related to C9 Aggrephagy Occur *In Vivo*

We examined the relevance of C9 aggregates and attendant inflammatory signaling *in vivo.* We initially asked whether C9 aggregates could occur in AV. To do this, we initially employed a humanized model of ischemia reperfusion injury (IRI).^25^ In this model, human artery segments are subjected to hypoxia in organ culture prior to being implanted as interposition xenografts in descending aortae of SCID/beige immunodeficient mice pre-engrafted with human lymphoid cells. We previously found that this protocol activates complement (C’) and forms MACs in ECs but also the media,^25^ a region relatively spared from collagen deposition in AV.^36^ This allowed a region for assessing thioflavin staining without confounding collagen fluorescence. Compared to controls, complement-treated grafts developing AV showed increased thioflavin staining in medial regions (Figure 8*a*), and complement-treated tissues showed increased insoluble C9 (Figure 8*b*). This indicated that C9 aggregates could be induced in human tissues *in vivo*.

**Figure 8.**
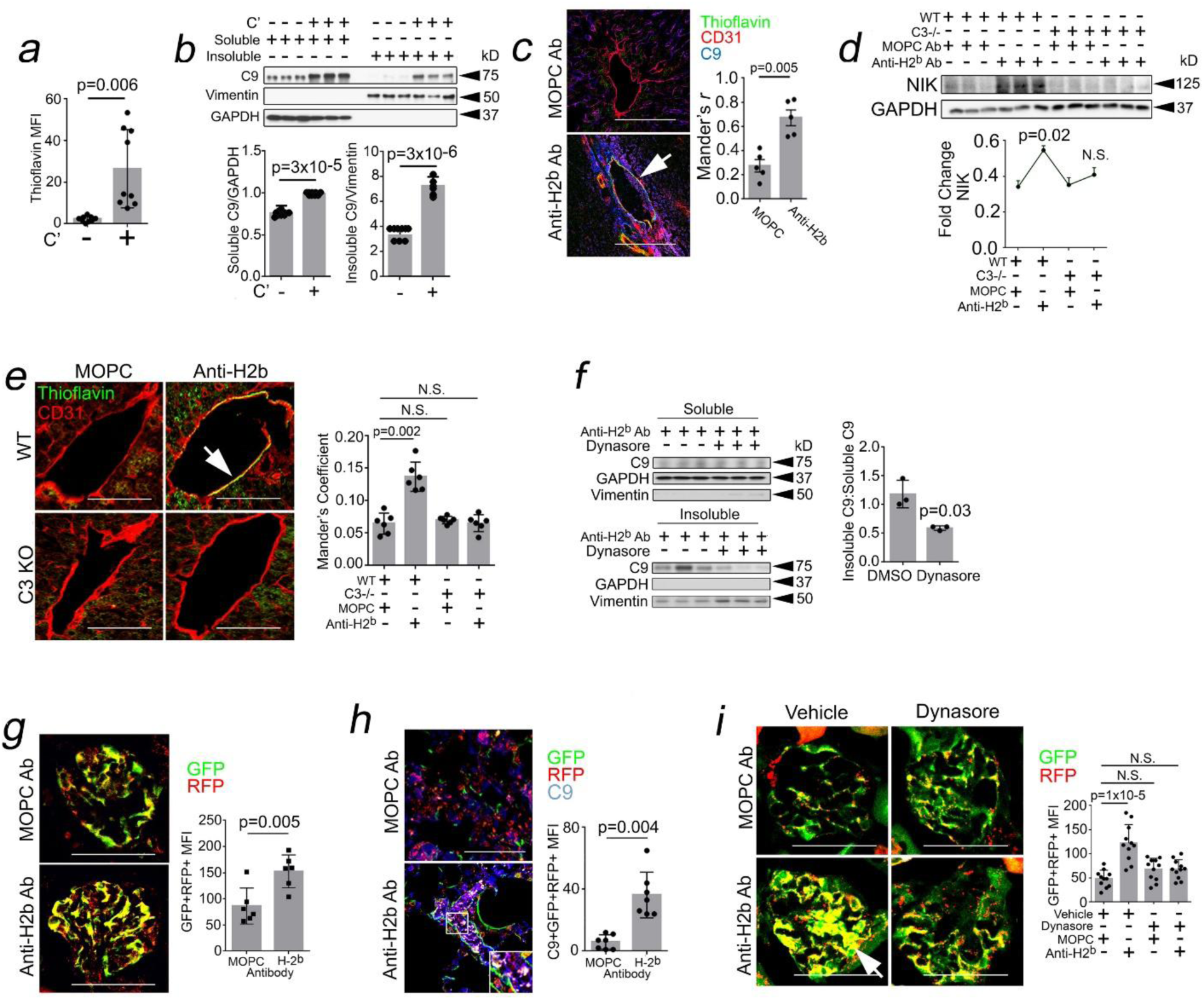
Signaling Responses Related to C9 Aggrephagy Occur *In Vivo.* Human coronary arteries were subjected to normoxia or hypoxia to induce complement (C’) activation. Arteries were subsequently implanted as xenografts into SCID/beige hosts engrafted with human lymphoid cells and analyzed 3 weeks post-implantation (*a,* n=8-9 per group). C’-treated human vessels were separated into soluble and insoluble lysate fractions and analyzed by Western blot (*b*). C57/Bl6 mice were injected with MOPC Ab (750μg) or anti-H2^b^ Ab (750μg), and liver tissues were harvested 24 hrs later for I.F. (*c*, n=5 per group). C3-/- mice were injected with MOPC Ab (750μg) or anti-H2^b^ Ab (750μg), and liver tissues were harvested 24 hrs later for Western blot (*d*, n=3 per group) and I.F. analysis (*e,* n=6 per group). C57/Bl6 mice were injected with Dynasore (100μg/mouse) once daily for 2 days prior to injection with anti-H2^b^ Ab, and the soluble and insoluble fractions of kidney lysates were analyzed 24 hours later (*f,* n=3 per group). Autophagy reporter (AR) mice were injected with MOPC Ab or anti-H2^b^ Ab and glomerular ECs were analyzed for GFP^+^RFP^+^ MFIs (*g*, n=6 per group) and C9^+^GFP^+^RFP^+^ MFIs (*h*, n=7 per group) 24 hrs later. AR mice were injected with Dynasore (100μg/mouse) once daily for 2 days prior to injection with MOPC Ab or anti-H2^b^ Ab, and glomeruli were analyzed 24 hours later (*i,* n=11 per group). Data points indicate individual murine hosts with >5 hpfs analyzed and averaged per host (*a-i*). Error bars indicate standard deviations. Scale Bar = 40μm (*c,e*), 10μm (*h*), and 20μm (*g*,*i*). One-way ANOVA with Tukey’s post-hoc correction (*d-e,i*) or paired Student’s *t*-test (*a-c, f-h*).

Inbred mouse strains express codified class I MHC regions called haplotypes, and C57/Bl6 mice express the H-2^b^ haplotype. To consolidate findings, we employed a second approach where we injected C57/Bl6 mice expressing the H-2^b^ haplotype with anti-H2^b^ Ab, a treatment we previously found could form non-cytolytic MACs on ECs following i.v. injection.^23^ With anti-H2^b^ Ab but not control MOPC Ab, C9^+^ ECs co-staining for thioflavin were visualized (Figure 8*c*), and anti-H2^b^ Ab-treated hosts showed increased NIK, a marker of NF-κB (Figure 8*d*). We found that NIK became significantly attenuated in C3^-/-^ mice (H-2^b^) whose ECs are capable of binding IgG but lack the ability to form MACs (Figure 8*d*). Concomitant with loss of NF-κB activity, C3^-/-^ mice showed decreased thioflavin staining in ECs (Figure 8*e*), indicating that C’ activation is required for endothelial aggregates and EC activation *in vivo*. We further observed that insoluble C9 aggregates were significantly reduced by Dynasore (Figure 8*f*), a CME inhibitor, indicating that *in vivo*, similar to PRA-treated HUVECs, C9 aggregates formed intracellularly. Autophagy reporter (AR) knockin mice express mRFP-GFP-LC3B, allowing autophagic flux to be assessed via GFP^+^RFP^+^ MFIs. We injected AR mice with anti-H2^b^ Ab and observed increased GFP^+^RFP^+^ regions (Figure 8*g*) colocalizing with C9 (Figure 8*h*), indicating increased C9^+^ autophagosomes. Notably, we observed that GFP^+^RFP^+^ regions were reduced by Dynasore (Figure 8*i*). Internalization of terminal complement proteins induces C9 aggregates, autophagic flux, and NF-κB activity *in vivo*.

### ZFYVE21 Mediates RNF34-Mediated Aggrephagy and NF-κB Activity *In Vivo*

Our data identified ZFYVE21 as a downstream effector employed by NUMBL to mediate C9 aggrephagy (Figure 7). To determine the relevance of ZFYVE2 signaling responses *in vivo,* in a first approach we generated ZFYVE21^fl/fl^ mice lacking exon 3 which in part encodes a FYVE domain required for ZFYVE21 colocalization to early endosomes.^50^ Using the breeding strategy in Figure 9*a*, we generated ZFYVE21^-/-^ mice globally lacking ZFYVE21. After confirming loss of ZFYVE21 in multiple tissues (Figure 9*b*), we assessed C9 aggregates following injection of ZFYVE21^-/-^ mice with anti-H2^b^ Ab. Relative to age- and sex-matched littermate controls, ZFYVE21^-/-^ mice showed increased thioflavin staining (Figure 9*c*) and increased insoluble C9 (Figure 9*d*), findings consistent with decreased C9 aggrephagy. These findings were associated with significantly reduced NF-κB activity marked by NIK and caspase-1 cleavage whose activity generates IL-1 family cytokines (lanes 4-6 *vs* lanes 10-12, Figure 9*e*). ZFYVE21^-/-^ kidneys treated with anti-H2^b^ Ab showed increased LC3B which, like P62, is known to become an autophagic substrate *in vivo,*^52^ indicating decreased autophagic flux (Figure 9*e*). Concurrent with the above, we found that MAC-treated hosts showed increased RNF34 (lanes 4-6 *vs* lanes 1-3, Figure 9*e*), and notably, RNF34 levels were virtually abolished in ZFYVE21^-/-^ mice under both basal and treatment conditions (Figure 9*e*), indicating a role for ZFYVE21 in mediating RNF34 stability *in vivo*.

**Figure 9.**
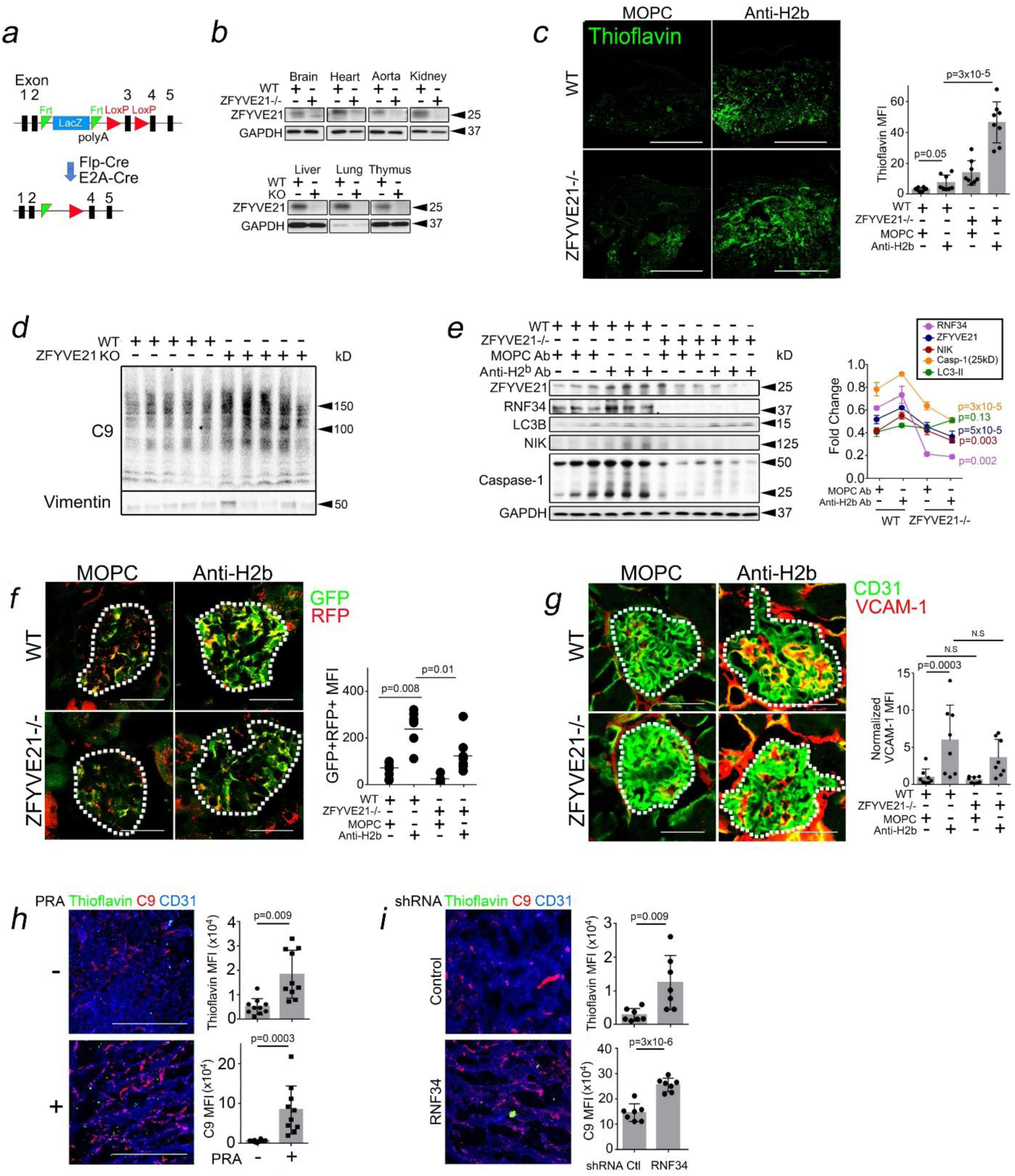
ZFYVE21 Mediates RNF34-Mediated Aggrephagy and NF-κB *In Vivo*. Generation of ZFYVE21^-/-^ mice (*a,b*). Six-week-old ZFVYVE21^-/-^ mice or littermate controls were injected with MOPC Ab or anti-H2^b^ Ab, and kidney tissues were analyzed for thioflavin by I.F. (*c,* n=8 per group). WT and ZFYVE21^-/-^ mice were treated with anti-H2^b^ Ab, and 24 hours later, insoluble fractions from kidney lysates were analyzed by Western blot (*d*, n=5 per group). WT and ZFYVE21^-/-^ mice were treated as indicated for 24 hours, and kidney lysates were analyzed by Western blot (*e,* n=3 per group). ZFYVE21^-/-^ x RFP-GFP-LC3B mice were treated with anti-H2^b^ Ab, and 24 hours later, glomerular GFP^+^RFP^+^ MFIs (*f*, n=5 per group) and CD31^+^VCAM1^+^ MFIs normalized to WT MOPC Ab-treated controls were assessed (*g,* n=7-9 per group). HUVECs were embedded in collagen-fibronectin gels and implanted subcutaneously into flanks of SCID/beige mice. Three weeks later, mice were intravenously injected with 200μL PRA, and gels were harvested and analyzed by I.F. 24 hours later (*h*, n=10 per group). HUVECs were transduced with shRNA as indicated prior to gel embedding and implantation into SCID/beige mice. Three weeks later, mice were intravenously injected with 200μL PRA, and gels were harvested and analyzed by I.F. 24 hours later (*i*, n=7 per group). Data points indicate total technical replicates for analyzed tissue regions (*c*) for n=3 hosts or n=7-10 hosts (*h,i*). Data points indicate biological replicates with n=5 (*f*) and n=7-9 (*g*). Error bars indicate standard deviations. Scale Bar = 75μm (*c*), 100μm (*f,g*) and 50μm (*h,i*). One-way ANOVA with Tukey’s post-hoc correction (*c,e,f,g*) or paired Student’s *t-*test (*h,i*).

In a second approach, we generated ZFYVE21^-/-^ x RFP-GFP-LC3B reporter mice, and we measured autophagic flux by quantifying GFP^+^RFP^+^ MFIs. Compared to controls showing robust autophagic flux with anti-H2^b^ Ab treatment, we found that ZFYVE21^-/-^ mice showed significantly reduced autophagic flux (Figure 9*f*). Tissue loss of ZFYVE21 and RNF34 in ZFYVE21^-/-^ mice was associated with reduced EC activation marked by VCAM-1 in glomerular ECs (Figure 9*g*).

In a third approach, we employed collagen-fibronectin gels containing HUVECs functionally lacking RNF34. HUVECs were embedded in collagen-fibronectin gels implanted subcutaneously into SCID/beige mice, a protocol allowing HUVECs to self-assemble into perfused microvessels *in vivo*.^23^ Three weeks post-implantation, hosts bearing collagen-fibronectin gels were treated with PRA. Twenty-four hours after PRA treatment, we observed Ulex^+^microvessels co-staining for C9, VCAM-1, and thioflavin (Supplementary Figure 6*c*). With PRA, gel-embedded microvessels additionally showed punctate C9^+^Thioflavin^+^ staining (Figure 9*h*). In follow-up studies, HUVECs were transduced with control or RNF34 shRNA prior to gel embedding and subcutaneous implantation. We found that RNF34 shRNA significantly potentiated both thioflavin and C9 MFIs in PRA-treated microvessels (Figure 9*i*).

To further assess the relevance of our *in vivo* findings, we analyzed public RNA seq data from complement-mediated conditions including CABMR (n = 110), rheumatoid arthritis (n = 23), and lupus nephritis (n = 21, Supplementary Figure 6*d*). These analyses showed significant correlations between genes marking EC activation and aggrephagy as well as moderate-high correlations between aggrephagy genes and both ZFYVE21 and RNF34. In 3 separate mouse models and in retrospective patient analyses, we find evidence that a ZFYVE21-RNF34 signaling axis mediates C9 aggrephagy, NF-κB, and EC activation *in vivo*.

### ZFYVE21 in ECs Dictates Alloimmune Tissue Injury

Finally, we examined immune effects of MACs including C9 aggregates *in vivo*. Male C57/Bl6 mice were injected with MOPC Ab or anti-H2^b^ Ab. The following day, Ab-treated skin grafts were placed onto C57/Bl6 mice containing lymphoid and myeloid cells, SCID/bg immunodeficient mice containing only myeloid cells, or SCID/bg mice pre-treated with liposomal clodronate (LP, 200μL) lacking both lymphoid and myeloid cells. After transplantation, allograft skin injury was assessed 21 days later by measuring epidermal thickening. While anti-H2^b^ Ab treatment increased epidermal thickening in all 3 treatment groups, epidermal thickening showed C57/Bl6 > SCID/bg >> SCID/bg + LC (Figure 10*a*). The C57/Bl6 and SCID/bg groups also consistently contained increased CD45^+^ cell infiltrates *vs* SCID/g + LC mice (Figure 10*b*). This indicated that MAC-induced tissue injury required contributions from both lymphoid and myeloid cells.

**Figure 10.**
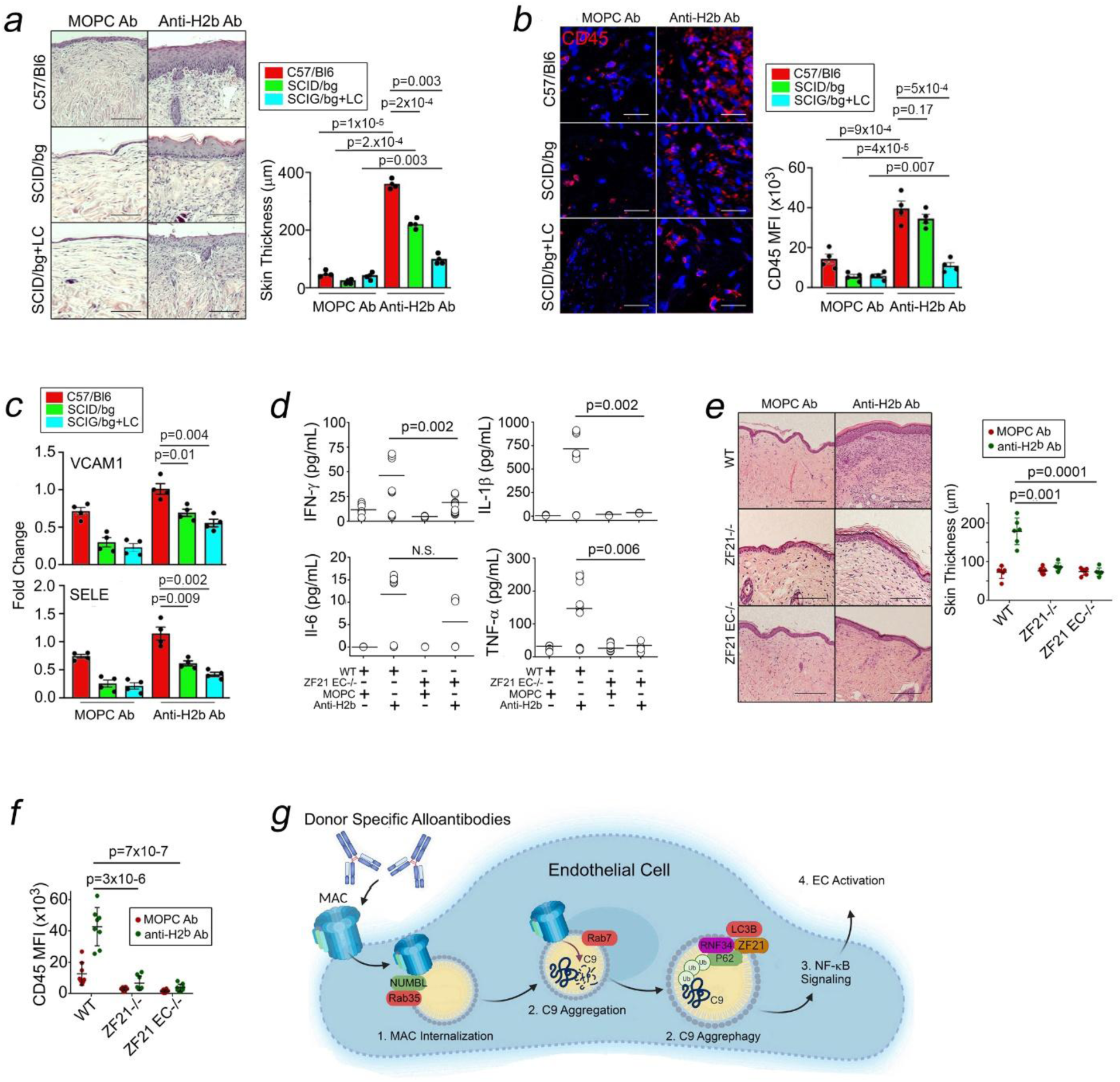
ZFYVE21 in ECs Dictates Alloimmune Tissue Injury. Male C57/Bl6 donors were treated with MOPC Ab (750μg) or anti-H2^b^ Ab (750μg), and 24 hrs later, Ab-treated skin were excised and placed as full-thickness allografts on female C57/Bl6, female SCID/bg, and female SCID/bg recipients pre-treated with 200μL liposomal clodronate i.p. 48hrs prior to skin grafting (SCID/bg + LC). Skin grafts were harvested 21 days later for H&E (*a*, n=4), I.F. (*b,* n=4), and qRT-PCR (*c*, n=4) and sera were analyzed for cytokines (*d*, n=12-17). Male WT, ZFYVE21^-/-^, and, ZFYVE21 EC^-/-^ donors were treated with MOPC or anti-H2^b^ Ab, and Ab-treated skin were placed on female SCID/bg recipients who then passively received 10x10^6^ female WT C57/Bl6 splenocytes intra-peritoneally. Skin tissues were harvested 21 days later for H&E (*e*, n=6) and I.F. analysis (*f,* n=8). Schematic showing how C9 aggregates may function as intracellular alarmins (*g*). Data points indicate biological replicates for skin regions (*a,b,e,f*), tissue samples (*c*), and sera (*d*). Error bars indicate standard deviations. Scale Bar = 100μm (*a,b,e,f*). One-way ANOVA (*c-f*) or two-way ANOVA (*a,b*) with Tukey’s post-hoc correction.

We next assessed relative contributions of lymphoid cells, myeloid cells, and EC intrinsic processes towards MAC-induced EC activation. In skin grafts above harvested from C57/Bl6, SCID/bg, and SCID/bg + LC mice, we assessed EC-specific VCAM and E-selectin (SELE) via qRT-PCR. We found that, in contrast to tissue injury requiring both lymphoid and myeloid contributions, EC activation showed C57/Bl6 > SCID/bg ≈ SCID/bg + LC (Figure 10*c*). Upon normalization, SCID/bg + LC mice showed ∼50% of the full transcript signals elicited in anti-H2^b^ Ab-treated C57/Bl6 hosts (Supplementary Figure 7*a*). This indicated that lymphoid cells and EC intrinsic processes dually contributed to the bulk of MAC-induced EC activation with myeloid cells playing relatively minor roles.

We next identified immune cell(s) activated by MAC-bound ECs. In a first approach using EC:PBMC cocultures, HUVECs were pre-treated with PRA to form C9 aggregates prior to co-culture with human PBMCs. After 10-14 day co-culture, we observed that MAC-treated HUVECs strongly enhanced both myeloid and lymphoid immune populations (Supplementary Figure 7*b*). We secondly confirmed *in vitro* findings above using a skin allograft model of CABMR.^26^ This model relies on the notion that MAC-bound donor ECs enhance recipient immune effector responses. Male skin allografts from C57/Bl6 mice were treated with anti-H2^b^ Ab and placed onto dorsal flanks of SCID/bg recipients prior to adoptive transfer of low numbers of female WT C57/Bl6 splenocytes responding to the minor H-Y alloantigen. We previously showed that this model enables both alloAb-induced MAC assembly and tissue pathologies consistent with chronic vascular inflammation.^23^ At 21 days, we tested skin, blood, and splenic immune cells. Consistent with EC:PBMC cocultures, MAC-bound ECs contributed to infiltration (lane 1 *vs* lane 3, Supplementary Figure 7*c*) and effector function (lane 1 *vs* lane 3, Supplementary Figure 7*d*) of both myeloid and lymphoid cells within all 3 tissue compartments tested. These data indicated that MAC-bound ECs mediated systemic effects, expanding both lymphoid and myeloid populations. Compared to WT controls, Cdh5-Cre x ZFYVE21^fl/fl^ (ZF21 EC^-/-^) mice showed consistently reduced lymphoid responses in the skin compartment ECs (red bars, lane 3 *vs* lane 4, Supplementary Figure 7*d*), causing significantly reduced inflammatory cytokines (Figure 10*d*) and chemokines (Supplementary Figure 7*e*) in sera. This indicated a role for EC-specific ZFYVE21 for potentiating local T cell responses where MAC deposition occurred.

We next compared effects of EC- *vs* non-EC-derived ZFYVE21 on immune cell activation, using littermate control (WT), ZF21^-/-^, and ZF21 EC^-/-^ skin grafts. WT skin grafts showed increased epidermal thickening (Figure 10*e*) and CD45^+^ infiltrates (Figure 10*f*), both of which became significantly reduced at nearly identical levels between ZF21^-/-^ and ZF21 EC^-/-^ skin grafts. This indicated that ZFYVE21 expression in ECs is a principal contributor to MAC-induced tissue injury. Original Western blot films are shown in Supplemental Figure 8-13. From these studies, we conclude that endothelial ZFYVE21 modulates MAC-induced tissue injury *in vivo*.

## DISCUSSION

Our data show that C9, a MAC protein, forms intracellular aggregates with intrinsic immunogenicity. We detected C9 aggregates in patient tissues, and we defined a non-cytolytic mechanism whereby C9 aggregates promote inflammatory signaling. (1) NUMBL is post-translationally stabilized on Rab35^+^ endosomes where it directly binds to and mediates internalization of C9; (2) internalized C9 enters endolysosomes whose intraluminal microenvironments promote C9 aggregates; and (3) C9 aggregates promote aggrephagy-dependent NF-κB, leading to EC activation. For aggrephagy to occur, C9 aggregates are ubiquitinylated by RNF34, ubiquitinylated C9 binds P62, and C9-RNF34-P62 complexes are bridged to LC3B^+^ aggresomes by ZFYVE21, which we identify as an LC3B-interacting protein (Figure 10*g*). We demonstrate the relevance of this sequence in mediating intracellular aggregates, NF-κB, and EC activation across various mouse models *in vivo*.

Our findings are consistent with behaviors reported for aggregate-prone proteins like amyloid precursor protein (APP)^56^ and prion protein (Prp).^57^ Like C9, these proteins basally localize at the cell surface and become immunogenic upon forming aggregates intracellularly. However, in contrast to these amyloidogenic proteins as well as intracellular complosomes,^58^ C9 is exogenously deposited on target ECs, and thus its overall protein burden may become dysregulated in a cell extrinsic manner. Our analyses involving patient specimens and cell cultures (Figure 1,2) suggest that conditions involving persistent and/or strongly dysregulated MAC assembly, as may occur with CABMR, RCC, and PRA treatment, favor C9 aggregates. Cell intrinsic perturbations, particularly those involving the endolysosomal pathway and/or intracellular complement signaling, might similarly enhance C9 aggregate formation, and such changes constitute an important and unexplored focus area.

Classically, alarmins connote intracellularly sequestered proteins like S100 or histones that are basally non-immunogenic but that acquire immunogenicity upon being externalized and sensed by recipient immune cells.^30^ Here, we alternatively posit that C9, a protein ‘sequestered’ in extracellular space where it is non-immunogenic, acquires alarmin-like properties when present intracellularly. In this paradigm, C9 is sensed, pathologically modified, and is capable of activating inflammatory signaling within the same cell. Prior to forming aggregates, intracellular C9 may exist in multiple bound states including intact MAC pores, soluble C9 monomers, or CD59-bound C5b-9 complexes. Future work defining which of these upstream source(s) contribute to C9 aggregates may yield insights for blocking pro-inflammatory effects associated with MACs.

Our studies elucidate certain protein complexes mediating trafficking of C9 aggregates including Rab35-NUMBL and ZFYVE21-RNF34-P62 complexes. These data further strengthen the burgeoning role of trafficking processes with inflammatory signals propagated by intracellular complement components. Future studies examining C9-associated trafficking may uncover new and potentially druggable mediators of EC activation. Our findings define a novel immune feature of MACs in addition to their widely studied cytotoxic effects.

## METHODS

### PRA Treatment

‘High’ panel reactive antibody (PRA) sera were obtained as pooled, de-identified sera from the tissue typing laboratory at Yale New Haven Hospital. ‘High’ PRA sera were taken from renal transplant candidates showing allo-sensitization of ≥80% and negative testing for numerous infectious agents.^24^ Prior to use, PRA sera were supplemented with human complement (Sigma, #S1764) at a ratio of 1 vial of lyophilized human complement per 25mL PRA sera. PRA sera supplemented as above can be used for 3 weeks, after which complement activity deteriorates. For PRA treatment, HUVEC were pre-treated with IFN-γ (50ng/mL, Invitrogen) for 48-72h prior to placement in gelatin veronal buffer (GVB, Sigma) at 25% v/v for the indicated times. In pulse-chase studies, HUVECs were pulsed with GVB containing 25% v/v PRA sera for 4h prior to washing and chased using GVB buffer alone for the indicated times. For treatment of human kidneys in organ culture, PRA was added at 1:4 ratio with GVB, for 6 hours prior to analysis.

PRA sera were fractionated into IgG^-^ and IgG^+^ fractions as previously described.^24^ Total IgG concentrations were first determined in intact sera prior to fractionation by ELISA (Invitrogen). Then, per manufacturer’s specifications using a MAbTrap Kit (GE Healthcare, Piscataway, NJ), 500μL of neat sera were diluted 1:1 in binding buffer and passed through the provided column pre-equilibrated with binding buffer. The column was washed with 5mL binding buffer to collect IgG^-^ fractions, and the IgG^+^ fraction was eluted using 5mL of elution buffer containing 650μL of neutralizing buffer. The 5mL volumes of IgG^+^ and IgG^-^ fractions were then serially concentrated and re-diluted in PBS using five 30-minute spins at 2100 x g in Amicon Ultra Centrifugal Filter Devices (EMD Millipore). All IgG^+^ fractions were brought to a final concentration equivalent to the total IgG concentration prior to sera fractionation. All isolated fractions were then used at 1:10 dilution in gelatin veronal buffer.

### HAP1 Cells

HAP1 cells were obtained from Sanquin (Amsterdam, The Netherlands) and include single or combinatorial knockouts of complement regulators, as previously described.^39^ For all experiments, CD46/CD55 double-knockout cells were used. Cells were cultured in Iscove’s Modified Dulbecco’s Medium (IMDM) supplemented with 10% fetal bovine serum (complete IMDM) at 37°C and 8% CO₂. Cultures were passaged 2–3 times per week upon reaching ∼80% confluency. Cells were routinely screened weekly for bacterial contamination and tested for mycoplasma every three months.

### Cell Viability Assay

Black, clear-bottom 96-well plates (Sigma, Japan; cat# CLS3603) were coated with 0.01% poly-L-lysine for 15 min at room temperature, dried for 1 h at 37°C, and stored at 4°C for up to one week. HAP1 cells were seeded at 30,000 cells per well in complete IMDM and allowed to adhere overnight at 37°C, 8% CO₂. Cells were then washed with DPBS and incubated for 4 h at 37°C, 8% CO₂ under the following conditions: 1% Tween-20, 10% NHS, 10% C5-depleted serum, or IMDM alone. An equal volume of IMDM containing 10 µM SYTOX Blue (final concentration 5 µM; ThermoFisher) was added, followed by a 15 min incubation at room temperature. Fluorescence was measured using a CLARIOstar plate reader (BMG, Germany). The experiment was repeated 3 times on different days, representing a biological triplicate. Background fluorescence from empty wells with IMDM and Sytox was subtracted, and cell death was normalized to the 1% Tween-20 condition.

### C9 Fluorescent Tagging

Purified human C9 (CompTech, USA) was labeled using the Alexa Fluor 568 Protein Labeling Kit (ThermoFisher, USA) according to the manufacturer’s instructions. Briefly, 500 µL of 1mg/mL C9 was mixed with 50 µL of 1 M sodium bicarbonate and incubated with Alexa Fluor 568 reactive ester for 1 h at room temperature followed by overnight incubation at 4°C with stirring. Labeled protein was separated from free dye by centrifugation at 1000 × g for 2 min using a Zeba Dye Removal column (ThermoFisher, USA) and eluted in Dulbecco’s phosphate-buffered saline (DPBS). Aliquots were stored at –80°C until use.

### Cryo-Light and Electron Microscopy grid preparation

Quantifoil™ R 2/2 200-mesh gold finder grids were glow-discharged in air and coated with 0.01% poly-L-lysine for 15 min at room temperature using custom 3D-printed holders compatible with 96-well plates.^59^ Grids were dried for 1 h at 37°C and stored at 4°C for up to one week. HAP1 cells were seeded at 35,000 cells per grid in complete IMDM, based on triplicate counts using a Countess cell counter (ThermoFisher, USA), and allowed to adhere for 3 h at 37°C, 8% CO₂. Media was replaced with fresh complete IMDM and incubated overnight. Grids were washed once with DPBS and incubated for 4 h at 37°C, 8% CO₂ in IMDM lacking FBS, containing 10% C9-depleted serum supplemented with 360 nM C9-AF568 (40%). Cells were then stained with LysoTracker™ Green DND-26 (500 nM), MitoTracker™ Deep Red (100 nM), and Hoechst 33342 (2 drops/mL; ThermoFisher, USA) for 15 min under the same conditions. After two DPBS washes, grids were maintained on ice prior to plunge freezing. Vitrification was performed using a Vitrobot Mark IV (ThermoFisher, USA) at 95% humidity and 21°C. Immediately before blotting, 3 µL PBS was added to the grid, followed by manual blotting for 15–20 s and plunging into liquid ethane. Samples were stored in liquid nitrogen until imaging.

### Cryo-SIM imaging and analysis

Grids were screened at beamline B24, Diamond Light Source, using a custom-built cryo-SIM system as previously described.^60^ Z-stacks were acquired in the green, red, and far-red channels.

### Cryo-Focused Ion Beam (FIB) lamella preparation and cryo-electron tomography data collection

Grids were clipped into autogrids (ThermoFisher, USA) and screened for ice thickness and cell confluency using a 200 kV Glacios electron microscope (ThermoFisher, USA). Grids were then transferred to an Aquilos 2 FIB/SEM ThermoFisher, USA) equipped with an integrated wide-field fluorescence microscope (iFLM) and a 175 L liquid nitrogen dewar (SubAngstrom). SEM tile sets were collected and aligned to Glacios atlas maps using three reference points in MAPS v3.34 (ThermoFisher, USA). Fluorescence Z-stacks (0.1 µm step size) were acquired across the grid and aligned accordingly. The iFLM setup included a 20× objective (Zeiss Epiplan-Apochromat, NA 0.7, piezo-driven), quadband filter cube (Semrock 556 LED-DA/FI/TR/Cy5-B-000), camera (Basler ace 2, Sony IMX541 CMOS sensor), and LED source (CoolLED; 365/450/550/635 nm). Acquisition settings were: 365 nm (3%, 120 ms), 450 nm (10%, 200 ms), 550 nm (1%, 300 ms), and 635 nm (10%, 40 ms). Milling sites were selected based on ice thickness, grid position, and the presence of Alexa Fluor 568 and LysoTracker Green signal, with no contribution from far-red or blue channels. Grids were sputter-coated with inorganic platinum for 15 s at 30 mA, followed by organometallic platinum deposition (Trimethyl (methylcyclopentadienyl) platinum (IV)) using the GIS for 45 s, and a second sputter coat for 15 s at 30 mA. Lamella positions, eucentric height, and milling angles were determined in AutoTEM 2.4 (ThermoFisher, USA). Milling was performed with a gallium ion beam at 30 kV, using stepwise currents of 1.0, 0.5, and 0.3 nA for coarse to fine milling. Final thinning was carried out at 50 and 30 pA, producing lamellae of ∼150–200 nm thickness. Z-stacks were acquired post-milling, followed by a final sputter coat (7 mA, 15 s). Grids were then transferred to a Titan Krios G2 TEM microscope (ThermoFisher, USA).

Search maps were acquired in Tomography v5.21, imported into MAPS, and aligned with iFLM Z-stacks to identify lamellae of interest based on colocalized LysoTracker Green and C9-AF568 signal, with no signal in blue or far-red channels (Supplementary Figure 2*d*). C9^+^LysoTracker^+^ were detected in 3 of 19 milled lamellae. Tilt series were collected using a dose-symmetric scheme (parameters in Supplementary Table S2). A total of 21 tomograms were acquired, including three control tomograms from lamellae lacking LysoTracker or C9 signal (Supplementary Figure 2*d*). Target areas included large electron-dense aggregates and associated vesicular networks corresponding to LysoTracker and C9-AF568 fluorescence.

### Tilt series alignment and reconstruction

Tilt series were processed using the WarpTools (v2.0.0) pipeline,^61^ including motion correction and CTF estimation, with handedness determination. Alignment was performed using the WarpTools IMOD (v5.1.7) patch-tracking wrapper, followed by manual refinement and curation in etomo.^62^ CTF-corrected tilt series were reconstructed into tomograms at a pixel size of 10 Å. Motion correction and reconstruction steps also generated half-tilt series to enable denoising using CryoCare (v0.3.0).^63^

### Cell Feature Segmentation

Ribosomes were automatically segmented using a pretrained model in EasyMode (v0.0.2, https://github.com/mgflast/easymode). Membranes were segmented using a pretrained model in MemBrain-v2 (v0.0.1).^64^ Aggregates were segmented using a trained neural network in the EMAN2-seg pipeline (v2.99.72)^65^ following the standard protocol. Tomograms were pre-processed using the EMAN2 GUI, including low-pass and high-pass filtering, pixel intensity normalization, and clamping of extreme values. The model was trained on manually selected good and bad particles (64 × 64 pixels), representing target features and background, respectively. Three tomograms from different lamellae and defocus values were used, with 20 good and ∼100 bad particle examples sampled across z-slices. Good particles were manually segmented using the paintbrush tool and used to train the network over 40 iterations. The model was iteratively refined through manual assessment, repicking, and retraining.

Ribosomes were automatically picked per tomogram using EasyMode based on the segmentation described above, giving ribosome frequency per tomogram. Aggregate volumes were quantified (in voxels) for each tomogram using the *imodauto* function in IMOD (v5.1.7).^62^ The lowest aggregate volume measured in a control tomogram from C9⁻ LysoTracker⁻ regions was used for background subtraction. Total tomogram volume was obtained from metadata, which was used to normalize the data. This was then expressed as a percentage of the total tomographic volume occupied by aggregates. To identify C1 in the tomograms, EMD-4232 was trimmed using the segmentation map tool in ChimeraX (v1.11.1)^66^ to remove density corresponding to the Fc region of bound IgG. The trimmed map was then manually fitted into the tomogram density using ArtiaX (v0.6.0)^67^ within ChimeraX. ^66^

### HUVEC Cell Culture Treatments

All protocols were approved by the Yale IRB (#25173). HUVECs were isolated as healthy, de-identified tissues from the Dept of Obstetrics and Gynecology at Yale New Haven Hospital as previously described and plated onto microtiter wells pre-coated with 1% gelatin.^21^ HEK293 cells were commercially obtained (ATCC). HUVEC were pooled from 3 human donors and cultured in complete EBM media (Lonza) containing bullet supplements (Lonza). Where indicated, HUVEC were pre-treated with Dynasore (80μM), Pitstop2 (30μM) for 30min, or bafilomycin (100nM) for 24h in GVB prior to PRA treatment. For studies involving NH_4_Cl, HUVEC were placed in NH_4_Cl (10μM) dissolved in GVB for 24 hrs in GVB prior to PRA treatment. Following indicated treatments, to assess cellular aggregates, thioflavin T (SigmaAldrich, #T1892), was added at final concentration of 10µM to HUVECs for 30min at 37°C prior to fluorescence measurements. Thioflavin T fluorescence was measured at an excitation wavelength of 450–480 nm and emission wavelength 520-550nm using ≥6 experimental replicates per treatment group (Molecular Devices, SpectraMax iD3).

CD4+CD45RA^-^ T cells were isolated from human PBMCs following depletion with anti-human CD45RA Ab (eBioscience, clone H100) and HLA-DR Ab (clone LB3.1, gift from Jack Strominger, Harvard University) using anti-human CD4 conjugated magnetic beads according to manufacturer’s specifications (Invitrogen) and co-cultured with allogeneic HUVEC pretreated with IFN-γ for 48 h and then pre-treated for an additional 6 hours with PRA sera, control sera or complement-inactivated PRA sera in round bottom 96-well tissue culture plates (BD Biosciences) at a T cell:EC ratio of 30:1 in a volume of 200 μL of RPMI 1640 containing 10% fetal bovine serum with 2% L-glutamine and 1% penicillin/streptomycin. T cell proliferation assayed by labeling T cells with CFSE at 5 μM (Invitrogen) and assessed by flow cytometry at seven days. All samples were acquired using a FACSCalibur flow cytometer (Becton Dickinson) and analyzed using FloJo computer software (TreeStar, Inc., Ashland, OR). Intracellular cytokine staining of activated T cells was also performed after 10-14 days of co-culture. To do so, PMA (50ng/mL) and ionomycin (500ng/mL, Sigma, St. Louis, MO) were added to culture medium 6 hours prior to staining and monensin (eBioscience, San Diego, CA) at 2μM was added 4 hours prior to fixation and permeabilization with Cytofix/Cytoperm staining per the manufacturer’s specifications (BD Biosciences). Permeabilized cells were stained using antibodies against IFN-γ (Biolegend, #502516) and IL-17 (eBioscience, #17-7179-42) at 1:50 dilution and analyzed using a FACSCalibur flow cytometer (BD Biosciences).

### HUVEC Lysate Treatments

For co-immunoprecipitations, cells were lysed in RIPA buffer without SDS (Cell Signaling) containing protease inhibitor tablets (Roche, 1 tablet per 10mL RIPA buffer) in 1.5mL Eppendorf tubes with gentle agitation for 1 hr at 4°C. Following this incubation, lysates were incubated with 1-3ug of antibody against Rab5 (Santa Cruz #sc,8008), ZFYVE21 (Novus Biologicals, #H00079038-B01P), HA-Tag (Bethyl,# A190-138A), or Ubiquitin (Thermofisher, # 13-1600) at 4^○^C overnight. The next day, samples were incubated with 25μL Protein A/G Magnetic Beads (Thermofisher, #88802) for 1.5 hr at room temperature and then were washed using TBS containing 3% Tween-20 three times prior to Western blotting as below. Whole cell lysates samples were harvested according to manuals of Pierce Protein A/G Magnetic Beads.

Following the above, 4X Laemli’s buffer (12μL) and 1mM DTT (6μL) were added to 32.5μL sample, and this mixture was heated for 95°C for 13min. Subsequently, samples were loaded onto pre-cast polyacrylamide gels (Bio-Rad), and proteins were electrophoretically separated and transferred to methanol-activated PVDF membranes at 4°C for 90 minutes. Membranes were washed for 15 minutes three times using Tris-buffered saline containing 0.1% Tween-20 pH 7.4 (TBS-T, AmericanBio), blocked with TBST containing 3% bovine serum albumin (Sigma) for 1 hr at room temperature, and incubated with primary antibody at 4^○^C overnight. Antibodies used for Western blotting were all used at 1:1000 dilution and included ZFYVE21 (Biorbyt, # orb221973), RNF34 (invitrogen, # PA5-113296), Rab5 (Santa Cruz Biotechnology, #sc-46692), Rab7 (Cell Signaling, #9367), NUMBL (Bethyl, #A301-719A), K48-conjugated ubiquitin (Cell Signaling, #8081), caspase-1 (Santa Cruz Biotechnology, #sc-392736), SQSTM1/p62 (Cell Signaling, #5114s), C9 (Abcam, # ab173302), ATG5 (Cell Signaling, # 12994s), ATG16L1 (Cell Signaling, # 8089s), LC3B (Cell Signaling, # 83506s), Flag Rabbit (Cell Signaling, # 2368s), Flag Mouse (Sigma, # F1804), RFP (Abcam, # ab62341), GFP Goat (Rockland, # 600101215) and ß-actin (Cell Signaling, # 4970S).

HUVECs were fractionated into soluble and insoluble fractions as previously described.^24^ For proteinase K treatments, soluble and insoluble C9 fractions were incubated with 12.5 μg/ml proteinase K for 30 min at RT prior to Western blot analysis. Proteostat staining (Enzo, #51023) was performing according to the manufacturer’s specifications.

### Immunofluorescence (IF) Staining

For immunostaining, HUVEC were grown on glass coverslips, fixed and permeabilized with ice cold methanol for 15min, and blocked with PBS containing 0.1% Tween and 5% FBS for 1hr at room temperature (PBST). Primary antibodies were then incubated overnight at 4C at 1:200 dilution using the following antibodies: Ulex (Vector Labs, #B-1065), p-P65 (Santa Cruz, #sc-8008), ZFYVE21 (Atlas, # HPA055721), NUMBL (Bethyl, #A301-719A), vimentin (Invitrogen, # MA1-19168), HSP70 (Invitrogen, # MA3-007), HA-Tag Rabbit (Cell Signaling, # 3724S), HA-Tag Rabbit (Cell Signaling, # 3724s), HA-Tag Mouse (Cell Signaling, # 2367s), Flag Rabbit (Cell Signaling, # 2368s), Flag Mouse (Sigma, # F1804). For thioflavin staining, slides were incubated with 5μM thioflavin T (Sigma, #596200) dissolved in PBS. The next morning slides were washed 3 times in PBS and incubated 2 hours at room temperature with secondary antibodies (Invitrogen, #A-31571, #A-31573, #A-31572, #A-21202, #A-11055, #A-21206, # A-31570, # A21447, # A-21432) at 1:200 dilution.

Following staining, slides were washed, air dried, and cover slipped using a DAPI mounting media (ImmunoGold with DAPI, Invitrogen). Immunofluorescence was visualized using a Leica SP8 confocal microscope.

For I.F. analyses of FFPE patient biopsies, sections were deparaffinized and hydrated. TUNEL staining was conducted according to the manufacturer’s specifications (Invitrogen, #C10619). Antigen retrieval was performed at 95°C for 1 h in Antigen Unmasking Solution (VectorLabs) and stained using antibodies against ZFYVE21 (Atlas), C5b-9 (Dako, # M0777) at 1:200 dilution at 4 °C overnight prior to addition of secondary Abs (1:500) and thioflavin T 5µM for 2 h at room temperature.

### siRNA Transfection of HUVECs

HUVEC were pre-treated with IFN-γ for 48 hours prior to siRNA transfection. siRNA targeting NUMBL, Rab35, ZFYVE21, ATG5, ATG16L1, DMN2 and RNF34 or non-targeting siRNA (target sequence UAA CGA CGC GAC GUA A) were purchased as pooled siRNA (Horizon Discovery) and transfected into HUVEC at ∼50-70% confluency in 24-well plates (BD Falcon). siRNAs were diluted at 20-40nM concentration in Opti-Mem culture media (Gibco) and mixed at equal volume with RNAiMax transfection reagent (Invitrogen) diluted 1:50 in Opti-Mem for 15 minutes at room temperature as per the manufacturer’s specifications. This mixture was then added to HUVEC cultures at 1:6 ratio for 37°C for 6 hours prior to washing and buffer exchange with EGM2. IFN-γ was added at 50ng/mL, and cells were then analyzed by Western blot, luciferase assay, RT-PCR, or T cell functional assays 48 hours later (72 hrs after transfection).

### Enzyme-linked immunosorbent assay (ELISA)

Purified human soluble C9 was purchased from Complement Technology. A Nunc 96-well flat-bottom MaxiSorp plate was coated with C9 protein (1μg/mL) as bait, diluted in pH 9.6 Na2CO3 buffer, overnight at 4°C. The plate was washed 3 times with PBS containing 0.05% Tween-20 (PBST) and blocked with PBST containing 3% BSA at room temperature for 1 hour. Subsequently, the plate was washed 3 times with PBST, then incubated overnight at 4°C with prey proteins, either BSA or His-tagged human NUMBL (MyBioSource, MBS7615688) diluted in blocking buffer. The next day, the plate was washed 6 times before overnight incubation at 4°C with primary antibodies directed against the prey proteins. The antibodies used were BSA (ProteinTech, 66201-1-Ig) and 6x-His Tag (Invitrogen, MA1-21315). The plate was washed 6 times with PBST and incubated with HRP-conjugated secondary antibodies for 1 hour at room temperature. After 8 washes with PBST, TMB was added as a substrate for 10 minutes, and the reaction was stopped with an equal volume of 2N H2SO4. The absorbance at 450nm was then measured.

### Real Time Quantitative Reverse Transcription-Polymerase Chain Reaction (Quantitative RT-PCR

RNA was isolated from treated HUVEC according to the manufacturer’s specifications (Qiagen) and reverse transcribed (Applied Biosystems, Foster City, CA). Respective cDNA was amplified in a CFX Realtime System (Biorad, Hercules, CA) at a volume of 20μL containing dilutions of 1:20 Taqman probe (Applied Biosystems), 1:2 Taqman Gene Expression Master Mix (Applied Biosystems), and 1:10 cDNA in ddH-2O. RT-PCR gene probes were purchased from Applied Biosystems [IL6 (#Hs00985639_m1), SELE (#Hs00950401_m1)]. For amplification, samples were heated to 50°C for 2 minutes for one cycle, 95°C for 10 minutes for one cycle, and then 40 cycles where samples were heated to 95°C for 15 seconds and cooled at 60°C for 1 minute.

### Cell-Free Studies

In cell-free experiments involving complement proteins, human complement C9 (Complement Technology, #A126) was incubated at 0.125μg/μL in 50mM NaCl titrated to the pH indicated in the text and containing thioflavin (5μM) at a final volume of 50μL/well and gently rocked at 37°C for the times indicated prior to assessing fluorescence at excitation wavelength at 349nm and emission wavelength at 454nm using ≥6 experimental replicates per treatment group. In other studies, human complement proteins (C5 #A120, C6 #A123, C7, #A124, C8, #A125, and C9, #A126, Complement Technology) were incubated at 0.2μg/μL in a hypotonic buffer (50mM NaCl) titrated to pH 4.5 as indicated in the text. In cell-free experiments involving ubiquitin (Ub), human ZFYVE21 (0.4 μg), C9 (0.4 μg), P62 (0.4 μg), and Ub (1.0 μg) were co-incubated at a final volume of 50μL with gentle rocking at 4°C overnight prior to performing co-immunoprecipitations using an anti-Ub mAb (Invitrogen, #13-1600).

### Time-Lapse Imaging

For time-lapse imaging, colocalization analysis was performed using the ImageJ plugin Coloc 2. Prior to analysis, raw 16-bit two-channel image stacks were preprocessed in ImageJ as follows. A Gaussian blur (σ = 1) was first applied to all slices to reduce image noise, after which the images were converted from 16-bit to 8-bit format. The channels were then separated into individual stacks, and background subtraction was performed independently for each channel. A second Gaussian blur (σ = 0.8) was subsequently applied to each channel to further smooth the signal. The same preprocessing parameters were applied to all image stacks. For C9+ZFYVE21+ vesicle analysis, an overlapping-signal image stack was generated using the ImageJ Image Calculator with the "AND" operation. The size of individual vesicles at each recorded time point was measured using the Analyze Particles function based on binarized vesicle images, and vesicle displacement was quantified using the TrackMate plugin in ImageJ.

### Molecular Cloning Studies

NF-κB luciferase lentiviral particles were commercially obtained (Cignal Reporter Assay, Qiagen) and used at an MOI of 20 to infect HUVECs for 8 hrs two times as previously described.^23^ The following reporter constructs were obtained from Addgene: mCherry-Rab5 WT (a gift from Gia Voeltz, plasmid #49201), mCherry-Rab5 DN (a gift from Sergio Grinstein, plasmid #35139), pRK5-HA-Ubiquitin-K48R (Addgene plasmid # 17604), pRK5-HA-Ubiquitin-WT (Addgene plasmid # 17608), HA-p62 (Addgene plasmid #28027), pcDNA3-GFP-LC3-RFP-LC3ΔG (Addgene plasmid, #168997), pmRFP-LC3(Addgene plasmid #21075).

### EC Coculture Studies

All protocols were approved by the Yale Institutional Review Board (#0601000969). PBMCs were isolated from leukopacks using density centrifugation as described previously and cryopreserved in liquid nitrogen.^24^ For EC:T cell cocultures, CD4^+^CD45RO^+^ T cells were isolated from thawed cryovials using magnetic bead separation kits (Miltenyi) with HLA-DR Ab (clone L243, Novus #NB100-77855) and CD45RA Ab negative depletion (10μL per cryovial, eBiosciences, 14-0458-82). For EC:T cell cocultures, HUVEC isolated from a single donor were grown in U-bottom 96-well microtiter plates, pretreated with human IFN-γ (50ng/mL, Invitrogen) for 48-72hr. On the day of the experiment, ECs were placed in gelatin veronal buffer containing 25% v/v of PRA sera for 6 hr prior to addition of human CD4^+^CD45RO^+^ T cells which were added at 0.5-1x106 cells/well at a volume of 200μL in RPMI (Gibco) supplemented with 5% FBS, 1.5% L-glutamine, and 1% penicillin/ streptomycin. After 10-14 days of co-culture in a humidified incubator at 5% CO2 and 37°C, T cells were harvested for FACS analysis.

For EC:PBMC cocultures, HUVEC isolated from a single donor were grown in U-bottom 96-well microtiter plates, pretreated with human IFN-γ (50ng/mL, Invitrogen) for 48-72hr. On the day of the experiment, ECs were placed in gelatin veronal buffer containing 25% v/v of PRA sera for 6 hr prior to addition of human PBMCs, which were added at 1x10^6^ cells/well at a volume of 200μL in RPMI (Gibco) supplemented with 5% FBS, 1.5% L-glutamine, and 1% penicillin/ streptomycin. After 10-14 days of co-culture in a humidified incubator at 5% CO2 and 37°C, PBMCs were harvested for FACS analysis.

### Mouse Studies

All protocols were approved by the Yale IACUC (#2023-20175). Human artery grafts were obtained from consenting heart transplant donors as de-identified samples at Yale New Haven Hospital. Donors were obtained from patients with non-ischemic cardiomyopathy (NICM) angiographically lacking pre-existing stenotic lesions. Human artery segments approximating the diameter of murine descending aortae were dissected and placed into organ culture in DMEM with 5% FBS at 0% FiO2 5% CO2 for 37°C for 8 hours prior to surgical reimplantation as interposition grafts into descending aortae of recipient female SCID/beige mice at 6-12-weeks of age, pre-engrafted with human lymphoid cells as previously described.^22^ Grafts were harvested three weeks later for I.F. analysis.

Adult mice aged 6-12 weeks old were used in our studies. Mice were housed in a standard animal maintenance facility under a 12-hour light/12-hour dark cycle. Wild-type C57/Bl6 mice (Jackson Labs, #000664), autophagy reporter mice (Jackson Labs, #027139), and C3^-/-^ mice (#029661, Jackson Labs) were injected i.v. via tail vein injection with 750μg anti-mouse MOPC Ab (Ichor, clone MPC-11) or 750μg anti-H-2^b^ Ab (Ichor, clone 8-24-3), and twenty-four hours later, kidneys were harvested for I.F. analyses and co-IP studies as indicated.

To generate global and EC-specific ZFYVE21^-/-^ mice, C57BL/6N-*A^tm1Brd^ Zfyve21^tm2a(EUCOMM)Wtsi/BayMmucd^* mice were initially purchased from Mutant Mouse Resource & Research Centers (catalog 043932-UCD). To generate this strain, the L1L2_Bact_P cassette was inserted at position 111823434 of chromosome 12 upstream of exon 3, a region we found was required for ZFYVE21 localization to early endosomes. Founder mice were crossed with B6N.129S4-Gt(ROSA)26Sortm1(FLP1)Dym/J (#016226, Jackson Labs) to generate *Zfyve21^fl/fl^* mice which were then crossed to either *Ella-Cre* mice (#003724, Jackson Labs) or *Cdh5-Cre* mice (#006137, Jackson Labs) to obtain global and EC-specific ZFYVE21^-/-^ mice, respectively, both of which express the H-2^b^ haplotype.

For skin graft experiments, skin from male C57/Bl6 mice at 6-12 weeks of age, ZFYVE21-deficient strains, and littermate controls were treated with MOPC Ab or anti-H-2^b^ Ab as above. Dorsal skin segments were harvested 24 hrs after Ab injection and implanted on the dorsal flanks of female SCID/bg hosts (#CBSCBG-F, Taconic) as full-thickness xenografts. Seven days later, mice were injected i.p. with 5x10^6^ female C57/Bl6 splenocytes, and skin grafts were harvested three weeks later for analysis.^23^ Epidermal thickness was quantified in 3-5 hpfs using morphometry, and mononuclear cell infiltration was estimated by 3 blinded observers.

### Proteomic Analyses

For proteomic analyses, HUVECs from 3 separate donors were stably transduced with ZFYVE21-GFP, grown to confluence in 15 T175 flasks, and treated with vehicle GVB (gelatin veronal buffer) or PRA 25% v/v for 45 minutes. Subsequently, cells were harvested and GFP pulldowns were performed according to the manufacturer’s specifications using GFP-Trap agarose beads (#gta, ProteinTech). Proteins were eluted from agarose beads and subjected to trypsin digestion and label-free proteomic analysis by mass spectrometry-liquid chromatography (MS-LC) as previously described.^22, 23^

### Multiplex Laser Bead Assay

Polystyrene beads containing fluorescent dyes were coated with capture antibody specific for a given protein analyte. Color-coded beads were then analyzed using a bead analyzer (Bio-Plex 200) containing a dual-laser system where the fluorescent dye within each bead is activated, and a second laser excites the fluorescent conjugate (streptavidin-PE) that has been bound to the beads during the assay. The amount of conjugate detected by the analyzer is in direct proportion to the amount of the target analyte which can be quantified using a standard curve (Eve Technologies).

### Statistical Methods & Software

Paired analyses were performed using paired Student’s t test and multiple comparisons were performed using a one-way or two-way ANOVA followed by Tukey’s pairwise comparison test using Origin computer software. p-values <0.05 were considered statistically significant. For segmentation analyses, data were tested for normality and then subjected to Welch’s *t*-test or Mann Whitney-U Test for the aggregate volume or number of ribosomes per tomogram, respectively. Standard deviations are reported throughout the text. The authors used an institutionally licensed copy of BioRender to produce Figure 10*g*.

## DATA AVAILABILITY

Source Data are provided with this paper. All data and methods are available from the authors upon reasonable request. The following transcriptomic datasets were retrieved from the Gene Expression Omnibus:

GSE147089-(n=224)-[https://www.ncbi.nlm.nih.gov/geo/query/acc.cgi?acc=GSE147089]

GSE112943-(n=21)-[https://www.ncbi.nlm.nih.gov/geo/query/acc.cgi?acc=GSE112943]

GSE97779-(n= 23)-[https://www.ncbi.nlm.nih.gov/geo/query/acc.cgi?acc=GSE97779].

For these datasets, an ‘Aggrephagy Index’ was calculated by averaging probe counts for 7 genes annotated to aggrephagy (ATG5, ATG16L, SQSTM1, ATG12, MAP1LC3B, BECN1, WDFY3), and an ‘EC Activation’ index was calculated by averaging probe counts for 3 adhesion molecules highly expressed in activated ECs (VCAM1, ICAM1, SELE).

## ACKNOWLEDGMENTS

We thank Diamond for access and support of the custom built cryoSIM beamline at B24 accessed through proposal BI40765. We thank the Structural Biology Science Technology Platform at the Crick and S. Islam at Imperial for technical support. All authors declare no financial conflicts of interest.

## AUTHOR CONTRIBUTIONS

G.S., Z.M., M.F., L.H., Q.J., M.N.B., performed Western blot experiments. Y.L., S.W., and Q.W. performed *in vivo* studies. G.S. and P.R. analyzed RNA-seq data. G.S., J.J., C.M., G.T., and G.M. performed I.F. analyses. J.C., Y.Z., and H.Z. performed public RNA-seq database analyses. X.G. and M.M. performed live cell imaging studies. D.P.N., D.B. and K.L.N performed cryoSIM experiments. D.P.N., A.N. and D.B. collected and analyzed cryo-FIB/SEM and cryo-ET data. G.S., Z.M., M.F., D.P.N., D.B., and D.J. designed experiments. G.S., Z.M., M.F., D.P.N., D.B., and D.J. analyzed experimental data. D.B. and D. J. drafted the manuscript.

