## Supplementary Information for "Internalized Components of Membrane Attack Complexes Disrupt Proteostasis and Acquire Alarmin-Like Properties"

**SUPPLEMENTARY TABLE**

**Supplementary Table S1. Patient Characteristics**

| **CABMR** | | | |
|---|---|---|---|
| **Age** | **Gender** | **Race** | **Transplant** |
| 27 | F | African-American | Kidney |
| 37 | M | African-American | Heart |
| 37 | M | African-American | Heart |
| 46 | M | Caucasian | Kidney |
| 55 | M | African-American | Heart |
| 56 | M | African-American | Kidney |
| 56 | F | Caucasian | Kidney |
| 57 | M | South Asian | Kidney |
| 60 | F | Caucasian | Kidney |
| 61 | F | Caucasian | Kidney |
| 65 | F | African-American | Kidney |
| 73 | F | Caucasian | Heart |
| **$C4d^+$ ABMR** | | | |
| 36 | F | Caucasian | Kidney |
| 42 | M | African-American | Kidney |
| 43 | M | African-American | Kidney |
| 52 | M | African-American | Kidney |
| **$C4d^-$ ABMR** | | | |
| 41 | M | Caucasian | Kidney |
| 47 | M | Caucasian | Kidney |
| 52 | M | African-American | Kidney |
| 56 | M | African-American | Kidney |
| **TCMR** | | | |
| 31 | M | African-American | Kidney |
| 42 | M | Caucasian | Kidney |
| 44 | M | Hispanic | Kidney |
| 46 | M | African-American | Kidney |
| **Age** | **Gender** | **Race** | **Transplant** |
| 27 | F | African-American | Kidney |
| 37 | M | African-American | Heart |
| 37 | M | African-American | Heart |
| 46 | M | Caucasian | Kidney |
| 55 | M | African-American | Heart |
| 56 | M | African-American | Kidney |
| 61 | F | African-American | Kidney |
| 73 | F | Caucasian | Heart |

**Supplementary S2. Data Acquisition Parameters for cryoEM Data Acquisition**

| Parameters | Positions 1 and 2 | Positions 3, 4 and 5 |
|---|---|---|
| Microscope | Titan Krios G2 | Titan Krios G2 |
| Voltage | 300 | 300 |
| Camera | Gatan K2 | Gatan K2 |
| Energy filter | Bioquantum | Bioquantum |
| Slit width (eV) | 20 | 20 |
| Magnification | 33000 | 33000 |
| Pixel Size (Å/pixel) | 4.33 | 4.33 |
| Does per tilt image (e/Å$^2$) | 3.2 | 3.2 |
| Total electron dose (e/Å$^2$) | 102.4 | 131 |
| Defocus range µm | 7 to 9 | 7 to 9 |
| Tilt range | -69 to 24 | -69 to 51 |
| Tilt increment | 3° | 3° |
| Software | Tomography 5.19 | Tomography 5.19 |

### SUPPLEMENTARY FIGURES

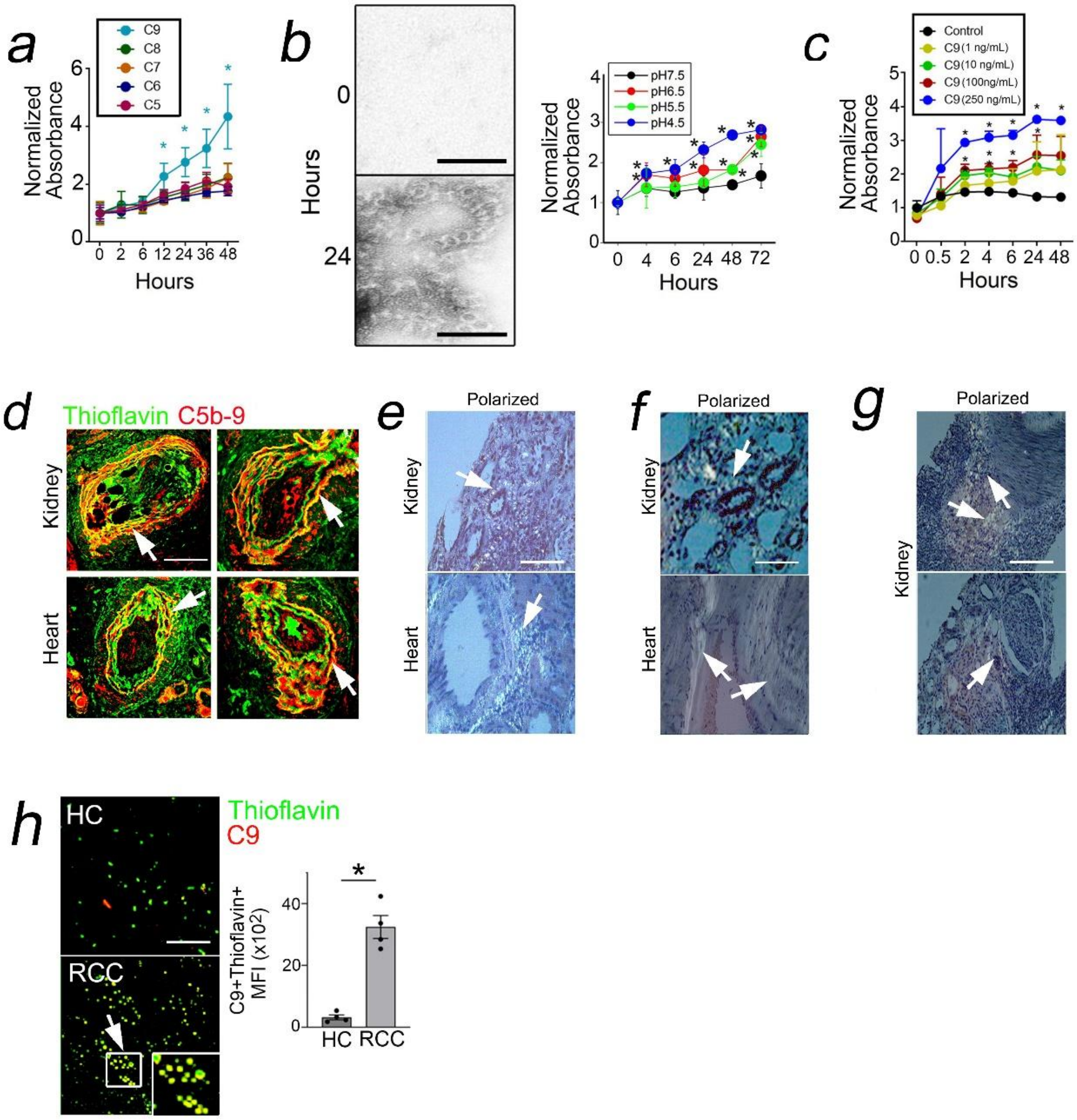


**Supplementary Figure 1.** Human complement proteins at 0.125 μg/mL were incubated at times indicated at pH 4.5 at 37°C (*a*). Human C9 protein (0.125 μg/mL) was resuspended in buffers of varying pH, and thioflavin fluorescence was assessed over time at 37°C (*b*). Human C9 protein at concentrations indicated were incubated at 37°C and thioflavin fluorescence was assessed over time (*c*). Human kidney and heart biopsies with CABMR were analyzed for thioflavin staining by I.F. (*d*, n=12). Kidney and heart biopsies with $C4d^-$ ABMR showing adventitial Congo red staining (*e*,*f*). Kidney biopsies with TCMR showing interstitial Congo Red staining (*g*). Kidney tissues from healthy controls (HC) and renal cell carcinoma (RCC) were analyzed by I.F. (*h*, n=4 per group). Experiments repeated ≥3 times using different HUVEC donors. Scale bars = 20nm (*b*), 200μm (*d*), and 100μm (*e-h*). Data points indicate technical replicates (*a-c*) or individual patient samples (*h*). * $p<0.05$ using two-way ANOVA (*a-c*) with Tukey's post-hoc correction or paired Student's *t*-test (*h*).

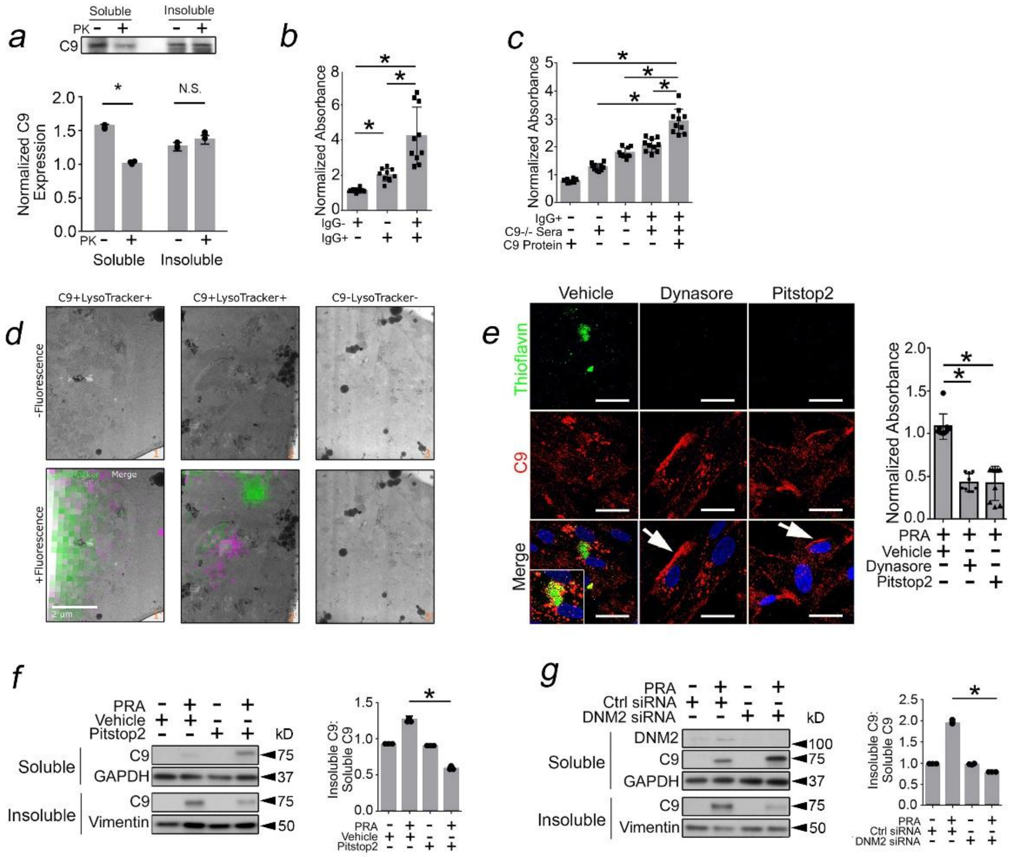


**Supplementary Figure 2.** SDS soluble and insoluble fractions of human C9 (0.125 μg/mL) were incubated at room temperature with proteinase K (12.5 μg/ml μg/mL) for 30 min prior to Western blot (*a*). C9 was normalized to GAPDH (lanes 1,2) or vimentin (lanes 3 and 4). PRA sera were fractionated and added to HUVECs for 2 hrs prior to assessing thioflavin fluorescence (*b*). $IgG^+$ fractions of PRA sera were added to HUVECs with $C9^{-/-}$ sera and C9 protein (5 μg/mL) for 2 hrs prior to assessing thioflavin fluorescence (*c*). Additional 2D images of milled lamellae used for cryoelectron tomography and for quantification, in addition to regions shown in Main Figure 2*h* (*d*). A negative-control field lacking C9 and LysoTracker staining is shown (*d*, right panel). The correlated fluorescence overlays show signal for both C9 (purple) and LysoTracker (green). Images 1 and 1′, 2 and 2′, and 3 and 3′ show the same fields of view without and with the fluorescence overlay, respectively. Fluorescence and electron microscopy modalities were aligned using three-point reference alignment in the MAPS software. HUVECs were pre-treated with vehicle, Dynasore (80 μg/mL), or Pitstop2 (30 μg/mL) for 30 min prior to addition of PRA for 4 hrs (*e*). Soluble and insoluble fractions of PRA-treated HUVECs exposed to vehicle or Pitstop2 were assessed for C9 (*f*). HUVECs transfected with control or DNM2 siRNA were treated with PRA, and soluble and insoluble lysates were assessed for C9 (*g*). Scale bars = 2μm (*d*) or 30μm (*e*). Experiments repeated ≥3 times using different HUVEC donors. Data points indicate technical replicates (*a-c,e-g*). * $p<0.05$ using one-way ANOVA (*e-g*) or two-way ANOVA (*b-c*) with Tukey's post-hoc correction or Student's *t*-test (*a*).

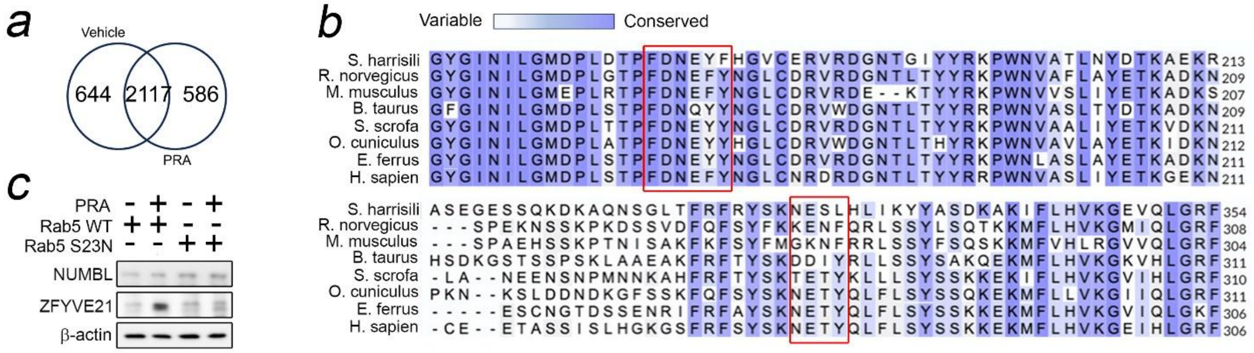


**Supplementary Figure 3.** ZFYVE21-GFP HUVECs were treated with PRA for 2 hours prior to GFP bead pulldowns and LC-MS/MS (*a,* n=3 runs). Sequence alignments predicted 2 NUMBL binding motifs on C9 (*b*). Western blots of HUVECs treated with PRA for 1 hour (*c*).

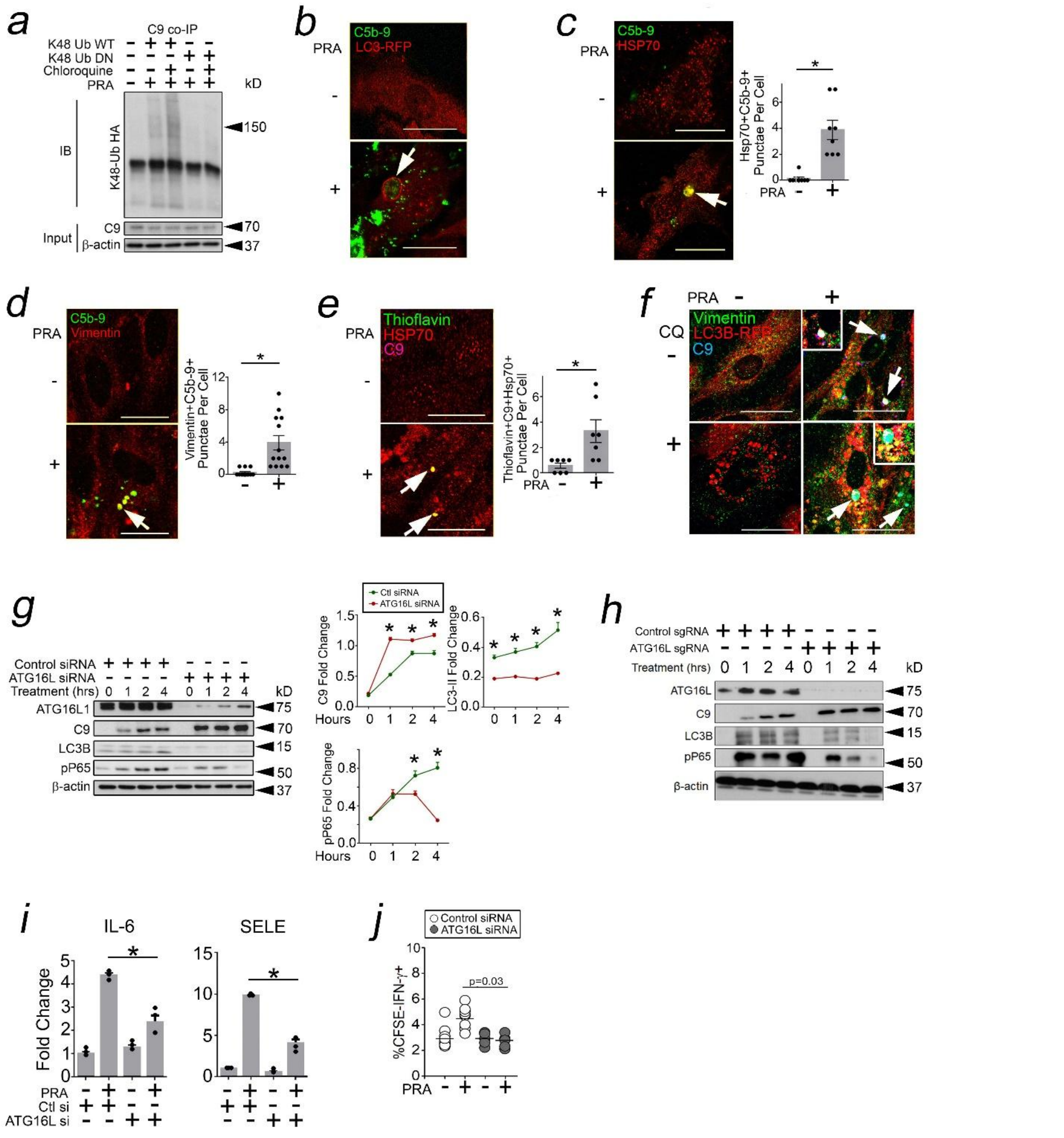


**Supplementary Figure 4.** K48 Ub-WT-HA was incubated with ZFYVE21, C9, and P62 proteins as indicated for 4 hours at 4°C prior to C9 pulldowns and Western blot analysis (*a*). HUVECs were treated with PRA for 2 hours prior to I.F. (*b-f*). HUVECs were transfected with ATG16L siRNA prior to pulse-chase studies with PRA (*g*). CRISPR/Cas9-mediated knockout of ATG16L was performed prior to pulse-chase studies with PRA (*h*). HUVECs were transfected with ATG16L siRNA prior to qRT-PCR (*i*), and EC:T cell cocultures (*j*) Data points indicate individual cells analyzed (*c-e*) or biological replicates (*i,j*). Scale bars = 30μm (*b-f*). * $p<0.05$ using one-way ANOVA (*i-j*) or two-way ANOVA (*g*) with Tukey's post-hoc correction or Student's *t*-test (*c-e*).

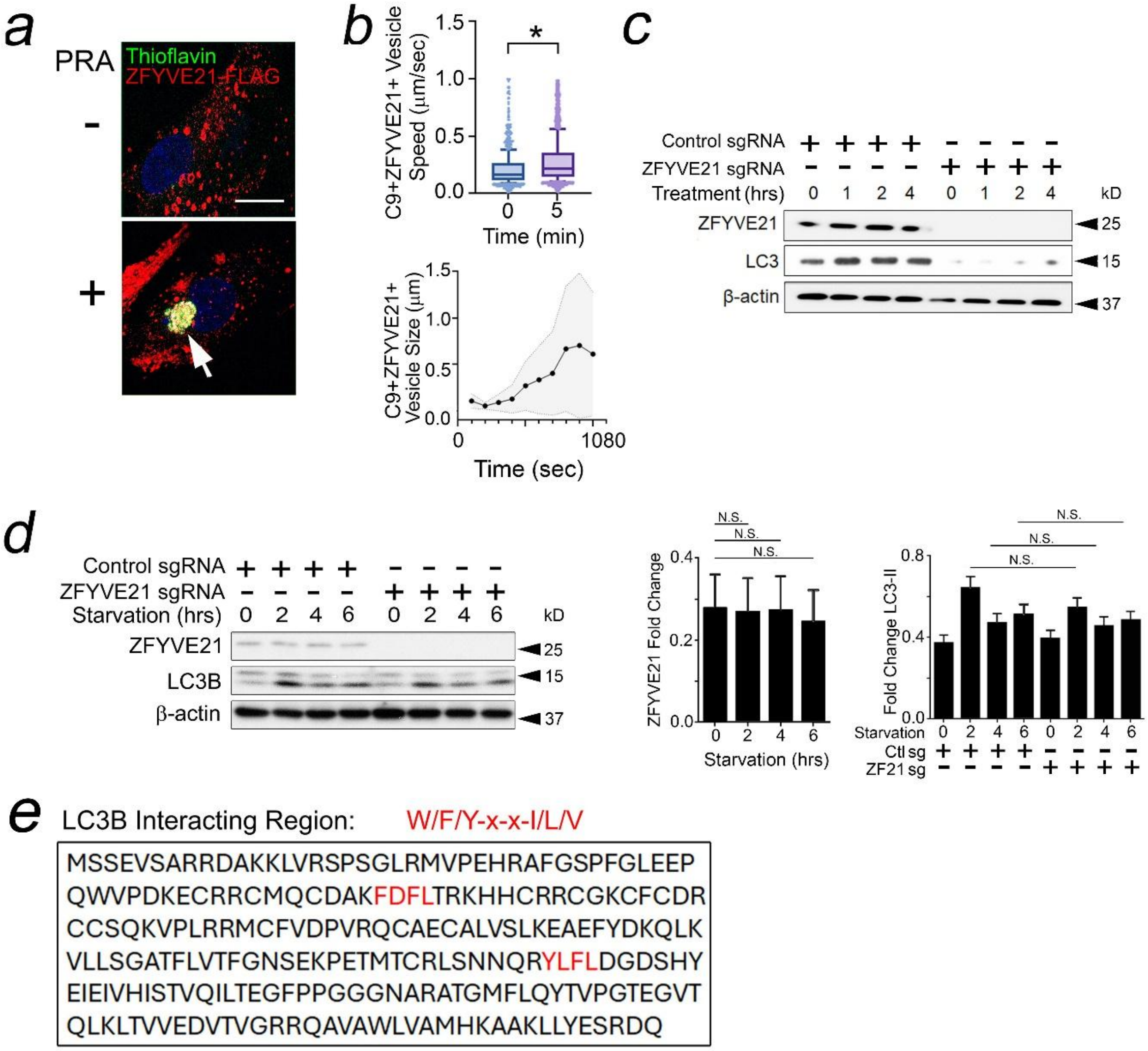


**Supplementary Figure 5.** HUVEC were stained as indicated after 2 hours of PRA treatment (*a*). Mean displacement (*b,* top) and mean size (bottom) for C9$^{+}$ZFYVE21$^{+}$ vesicles were calculated in live cell imaging studies. HUVECs were treated with PRA (*c*) or serum starved (*d*) as indicated prior to use in pulse-chase studies. Human ZFYVE21 protein sequence contains two LC3B interacting regions (*e*). Scale bar = 30μm (*a*). Data points indicate individual cells analyzed (*b*) or biological replicates (*c,d*). * $p<0.05$ using two-way ANOVA (*d*) with Tukey's post-hoc correction or Student's *t*-test (*b*).

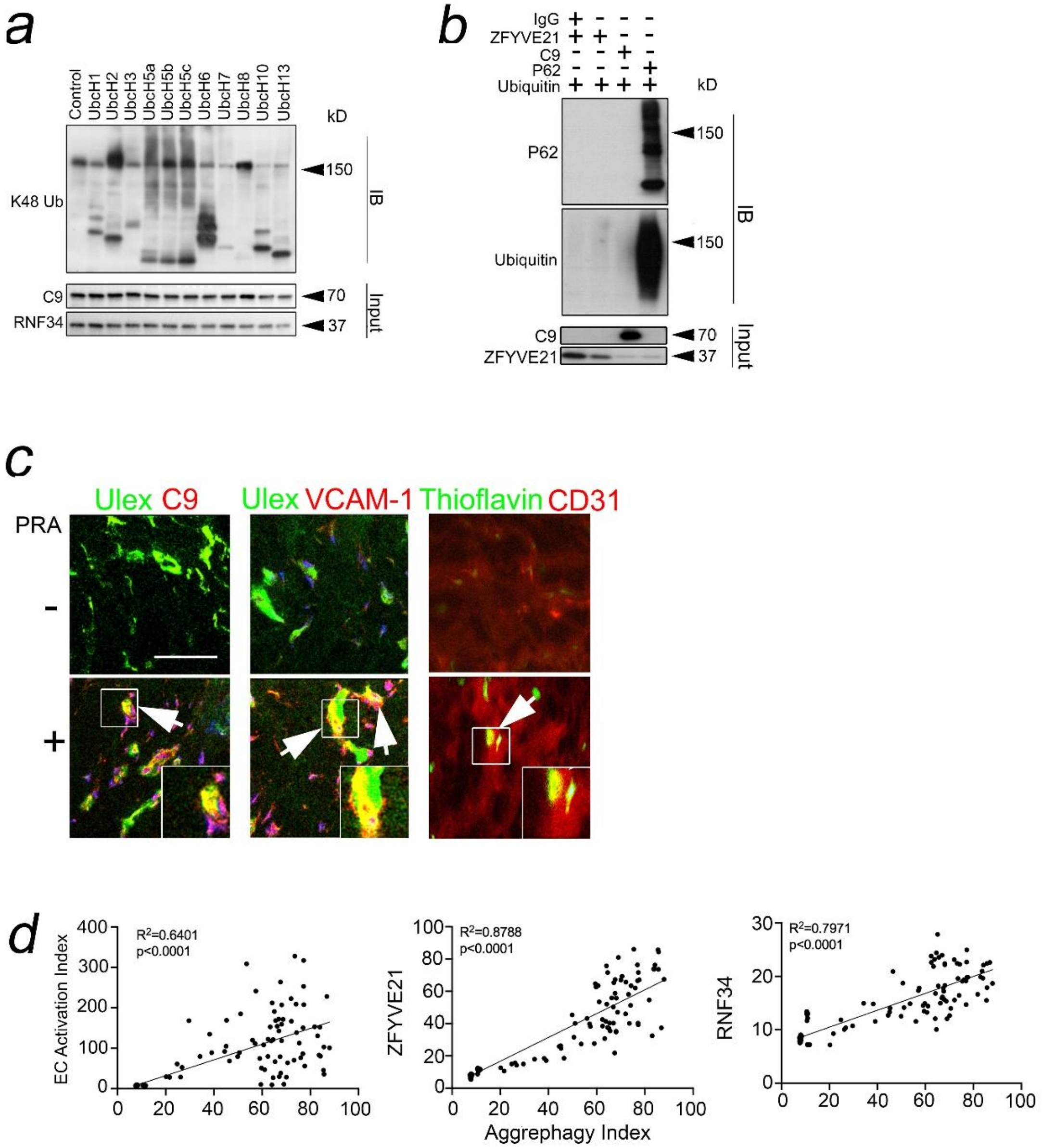


**Supplementary Figure 6.** Various E2 ubiquitin ligases were co-incubated with ubiquitin, E1 enzymes, RNF34 protein, and C9 protein and tested for *in vitro* ubiquitinylation of C9 for 4 hours at 37˚C (*a*). ZFYVE21, C9, or P62 proteins were incubated with Ub WT prior to Western blot analysis (*b*). HUVECs embedded in collagen-fibronectin gels and implanted in SCID/beige mice for 3 weeks prior to intravenous injection with 200μL neat PRA sera and gel harvest 24 hours later (*c,* n=3 per group). Correlation coefficients for genes indicated were calculated using public RNA-seq datasets (*d*). Scale bar = 20μm (*c*).

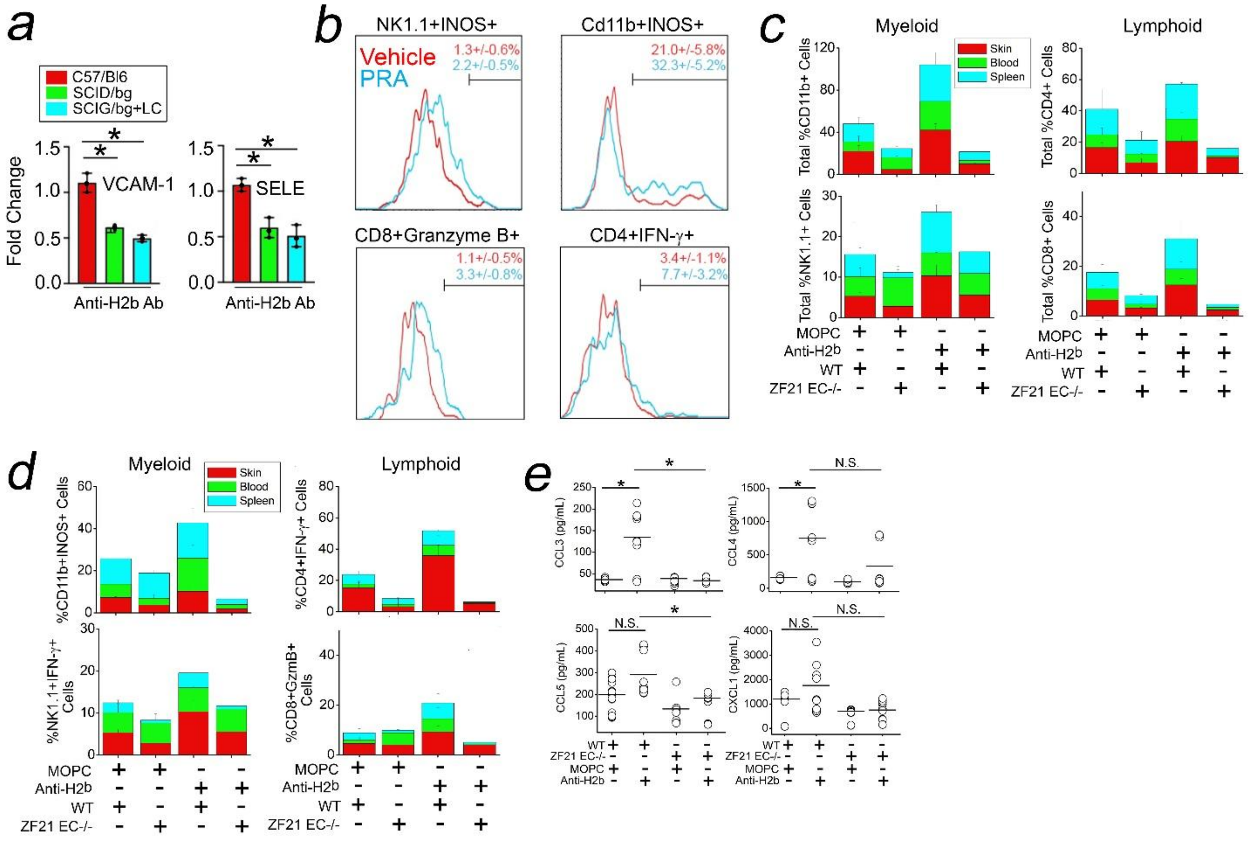


**Supplementary Figure 7.** qRT-PCR (*a*) and FACS analyses (*b*) were performed as indicated. Myeloid and lymphoid lineages expressed as a percent of total $CD45^+$ cells were analyzed in skin, blood, and splenic tissue compartments (*c,d*). Sera cytokine analysis (*e*). Data points indicate biological replicates (*a*, *e*). * p<0.05 using one-way ANOVA (*a*) or two-way ANOVA (*e*) with Tukey's post-hoc correction.

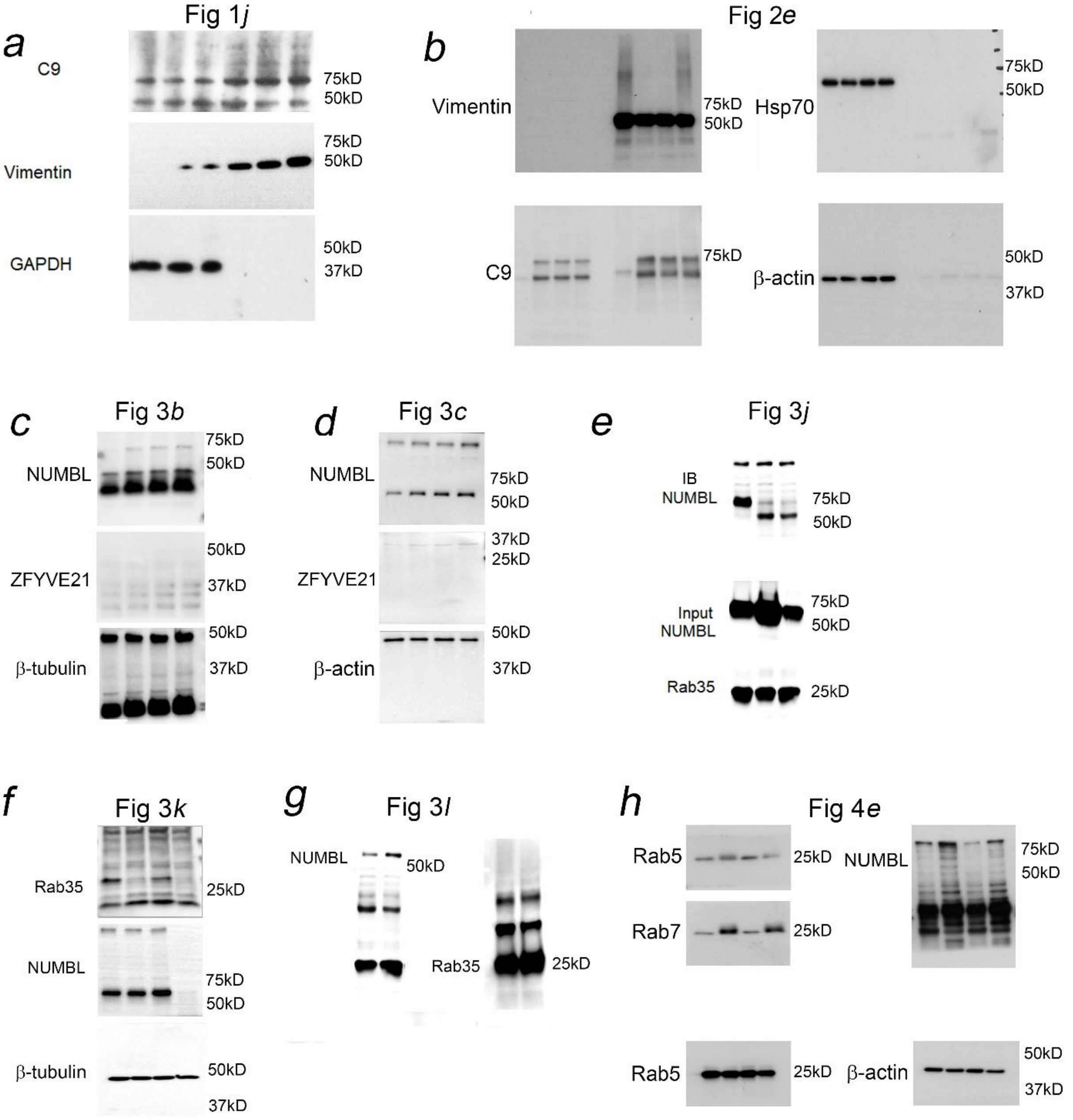


**Supplementary Figure 8.** Original uncropped films corresponding to Western blots in main Figures 1-4 in the manuscript.

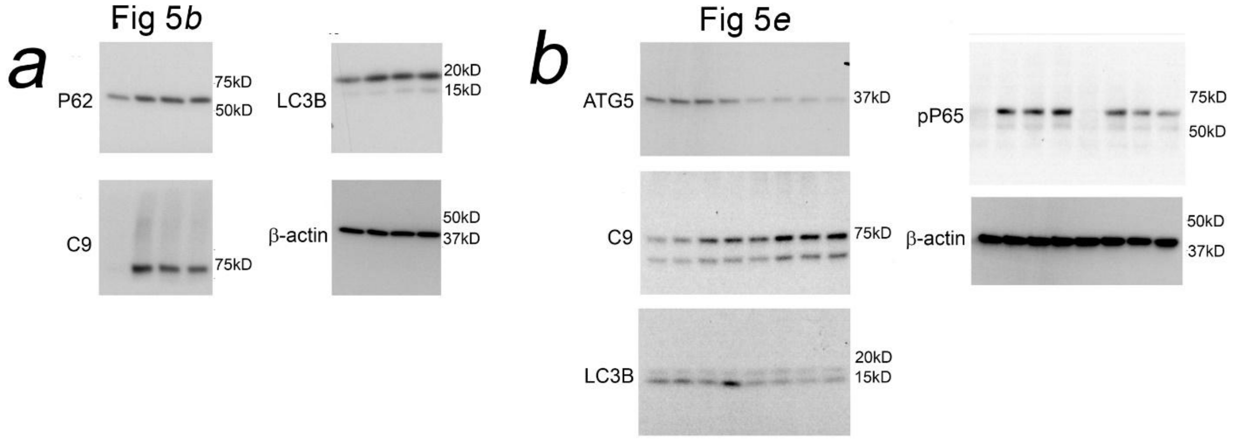


**Supplementary Figure 9.** Original uncropped films corresponding to Western blots in main Figure 5 in the manuscript.

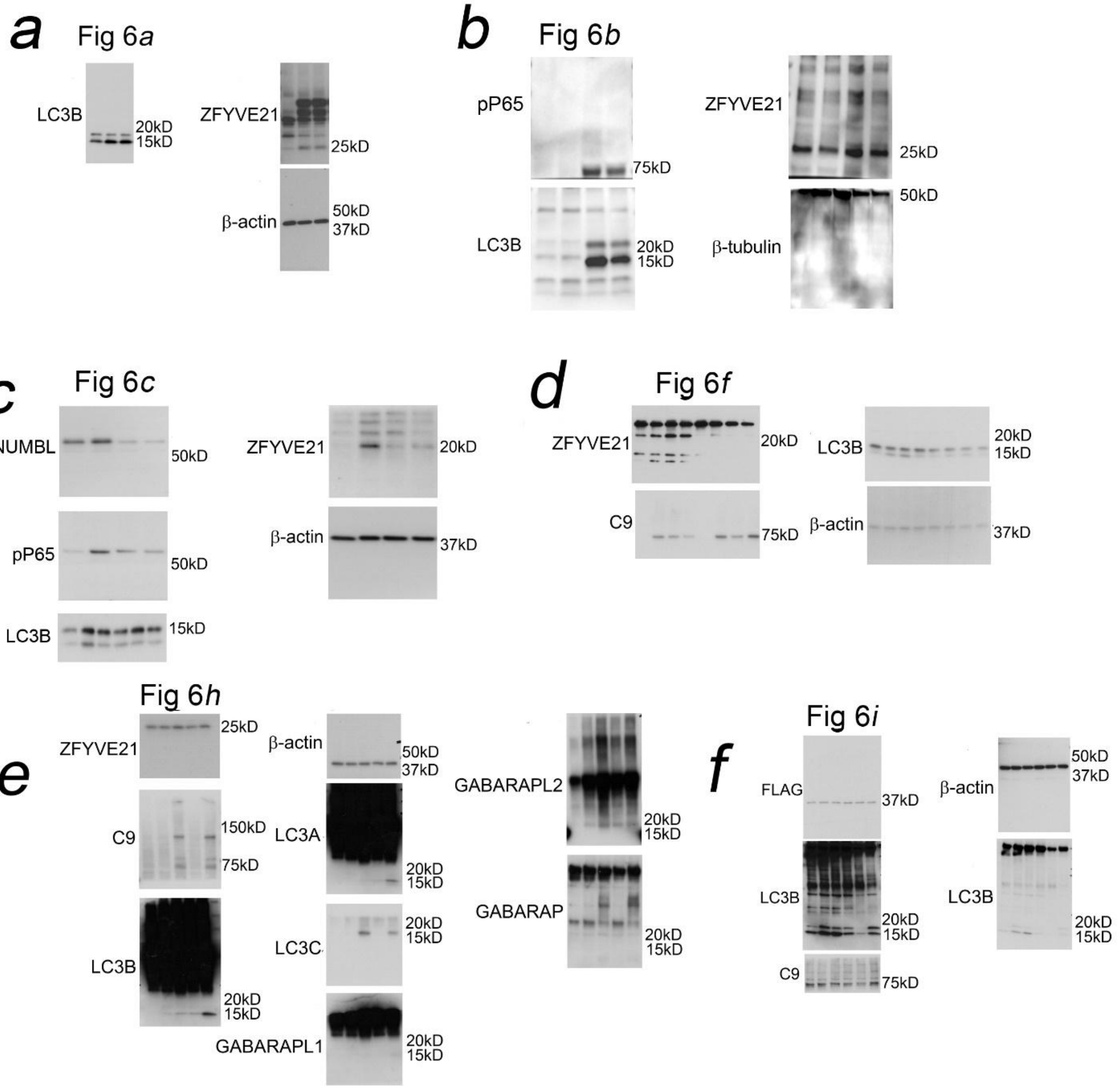


**Supplementary Figure 10.** Original uncropped films corresponding to Western blots in main Figure 6 in the manuscript.

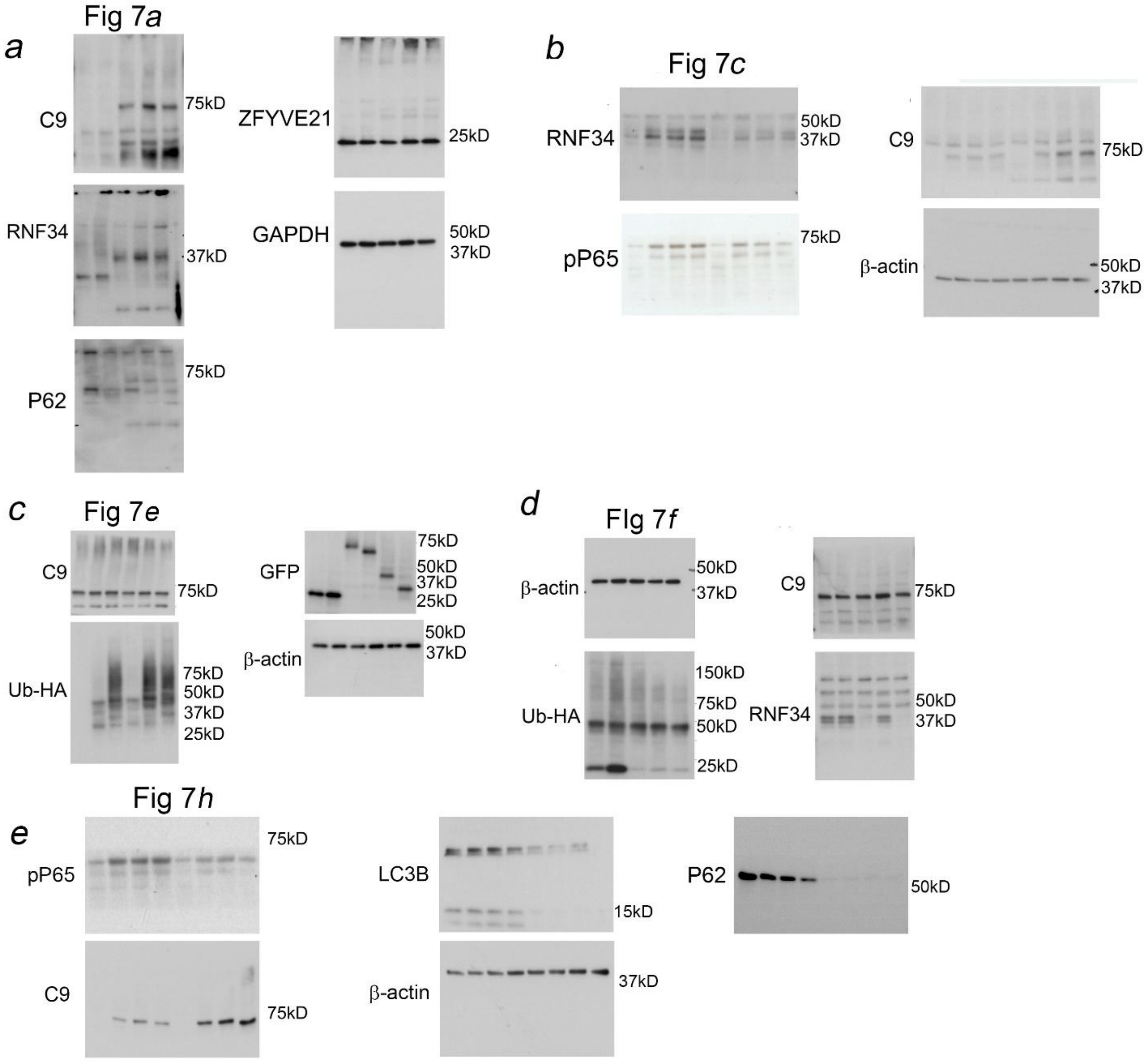


**Supplementary Figure 11.** Original uncropped films corresponding to Western blots in main Figure 7 in the manuscript.

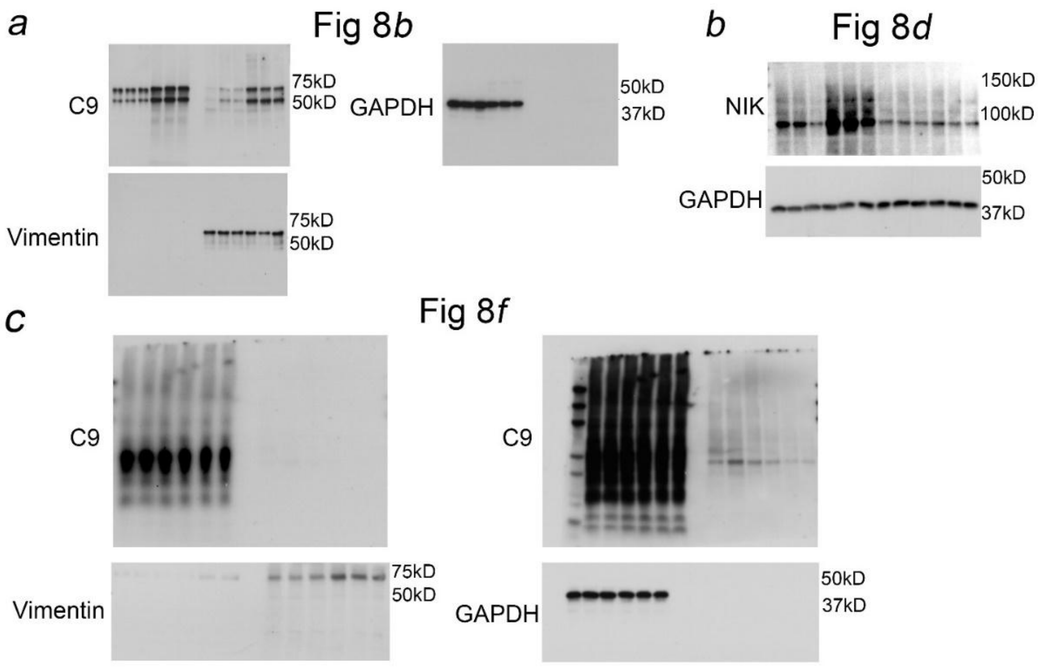


**Supplementary Figure 12.** Original uncropped films corresponding to Western blots in main Figure 8 in the manuscript.

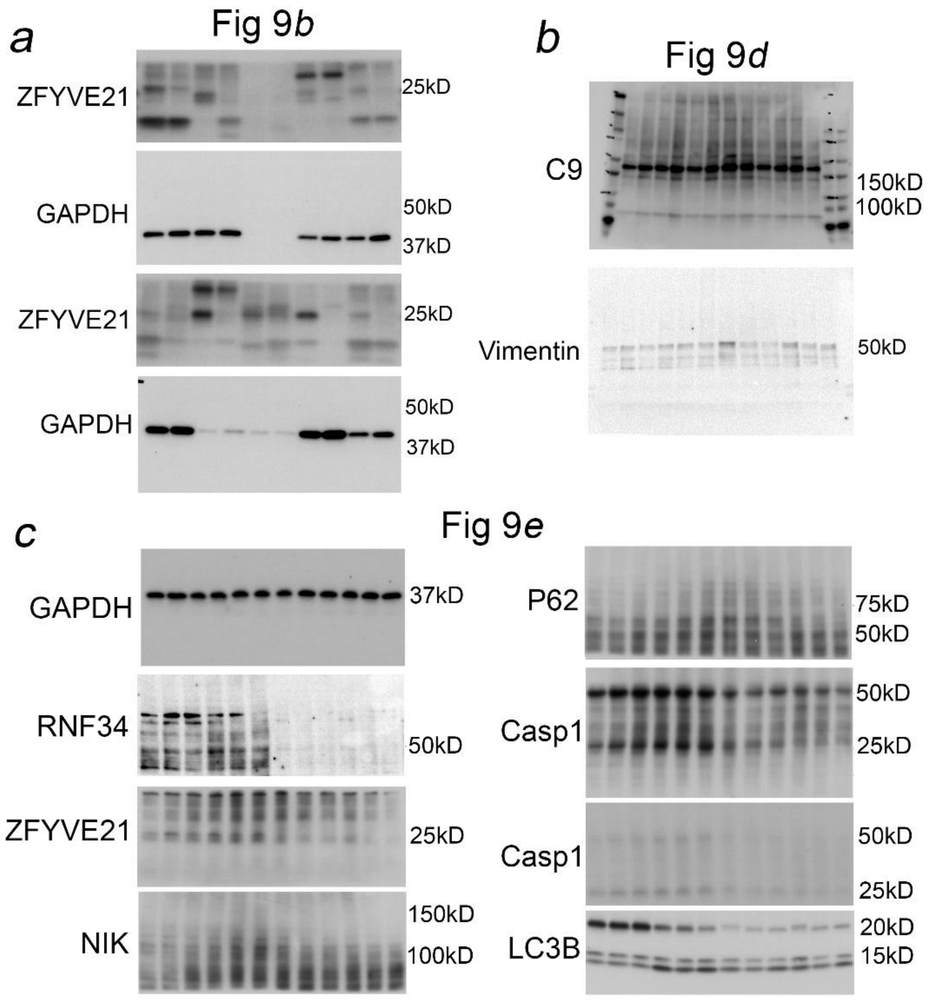


**Supplementary Figure 13.** Original uncropped films corresponding to Western blots in main Figure 9 in the manuscript.
